# Developmental function and state transitions of a gene expression oscillator in *C. elegans*

**DOI:** 10.1101/755421

**Authors:** Milou W.M. Meeuse, Yannick P. Hauser, Gert-Jan Hendriks, Jan Eglinger, Guy Bogaarts, Charisios Tsiairis, Helge Großhans

**Affiliations:** Friedrich Miescher Institute for Biomedical Research (FMI), Maulbeerstrasse 66, CH-4058 Basel; University of Basel, Petersplatz 1, CH-4001 Basel; University Hospital, Spitalstrasse 21, CH-4031 Basel

## Abstract

Gene expression oscillators can structure biological events temporally and spatially. Different biological functions benefit from distinct oscillator properties. Thus, finite developmental processes rely on oscillators that start and stop at specific times; a poorly understood behavior. Here, we have characterized a massive gene expression oscillator comprising >3,700 genes in *C. elegans* larvae. We report that oscillations initiate in embryos, arrest transiently after hatching and in response to perturbation, and cease in adults. Experimental observation of the transitions between oscillatory and non-oscillatory states at a resolution where we can identify bifurcation points reveals an oscillator operating near a Saddle Node on Invariant Cycle (SNIC) bifurcation. These findings constrain the architecture and mathematical models that can represent this oscillator. They also reveal that oscillator arrests occur reproducibly in a specific phase. Since we find oscillations to be coupled to developmental processes, including molting, this characteristic of SNIC bifurcations thus endows the oscillator with the potential to halt larval development at defined intervals, and thereby execute a developmental checkpoint function.

## Introduction

Gene expression oscillations occur in many biological systems as exemplified by circadian rhythms in metabolism and behavior (Panda et al., 2002), vertebrate somitogenesis (Oates et al., 2012), plant lateral root branching (Moreno-Risueno et al., 2010), and *C. elegans* larval development (Hendriks et al., 2014). They are well-suited for timekeeping, acting as molecular clocks that can provide a temporal, and thereby also spatial, structure for biological events (Uriu, 2016). This structure may represent external time, as illustrated by circadian clocks, or provide temporal organization of internal processes without direct reference to external time, as illustrated by somitogenesis clocks (Rensing et al., 2001).

Depending on these distinct functions, oscillators require different properties. Thus, robust representation of external time requires a stable period, i.e., the oscillator has to be compensated for variations in temperature and other environmental factors. It also benefits from a phase-resetting mechanism to permit moderate realignments, if needed, to external time. Intuitively, either feature seems unlikely to benefit developmental oscillators. By contrast, because developmental processes are finite, e.g., an organism has a characteristic number of somites, developmental oscillators need a start and an end. How such changes in oscillator activity occur *in vivo*, and which oscillator features enable them, is largely unknown (Riedel-Kruse et al., 2007; Shih et al., 2015).

Here, we characterize the recently discovered ‘*C. elegans* oscillator’ (Hendriks et al., 2014; Kim et al., 2013) at high temporal resolution and across the entire period of *C. elegans* development, from embryo to adult. The system is marked by a massive scale where ∼3,700 genes exhibit transcript level oscillations that are detectable, with large, stable amplitudes and widely dispersed expression peak times (i.e., peak phases), in lysates of whole animals. For the purpose of this study, and because insufficient information exists on the identities of core oscillator versus output genes, we define the entire system of oscillating genes as ‘the oscillator’. We demonstrate that the oscillations are coupled to molting, i.e., the cyclical process of new cuticle synthesis and old cuticle shedding that occurs at the end of each larval stage. We observe and characterize onset and offset of oscillations both during continuous development and upon perturbation, and find that these changes in the state of the oscillator system (or bifurcations), occur with a sudden change in amplitude. They also occur in a characteristic oscillator phase and thus at specific, recurring intervals. Because of the phase-locking of the oscillator and molting, arrests thus always occur at the same time during larval stages, around molt exit. This time coincides with the recurring windows of activity of a checkpoint that can halt larval development in response to nutritionally poor conditions. Hence, our results indicate that the *C. elegans* oscillator functions as a developmental clock whose architecture supports a developmental checkpoint function. Indeed, the features of the bifurcations constrain oscillator architecture, excluding a simple negative-loop design, and possible parameters of mathematical models.

## Results

### Thousands of genes with oscillatory expression during the four larval stages

Although previous reports agreed on the wide-spread occurrence of oscillatory gene expression in *C. elegans* larvae (Grün et al., 2014; Hendriks et al., 2014; Kim et al., 2013), the published data sets were either insufficiently temporally resolved or too short to characterize oscillations across *C. elegans* larval development. Hence, to understand the extent and features of these oscillations better, including their continuity throughout development, we performed two extended time course experiments to cover the entire period of post-embryonic development plus early adulthood at hourly resolution. We extracted total RNA from populations of animals synchronized by hatching in the absence of food. The first time course (designated TC1) covered the first 15 hours of development on food at 25°C, the second time course (TC2) covered the span of 5 hours through 48 hours after plating at 25°C. [Fig. S1A provides a summary of all sequencing time courses analyzed in this study.] The extensive overlap facilitated fusion of these two time courses into one long time course (TC3) (Fig. S1B), and a pairwise-correlation plot of gene expression over time showed periodic similarity that was repeated four times (Fig. 1A, light-gray off-diagonals), presumably reflecting progression through the four larval stages.

**Fig 1:**
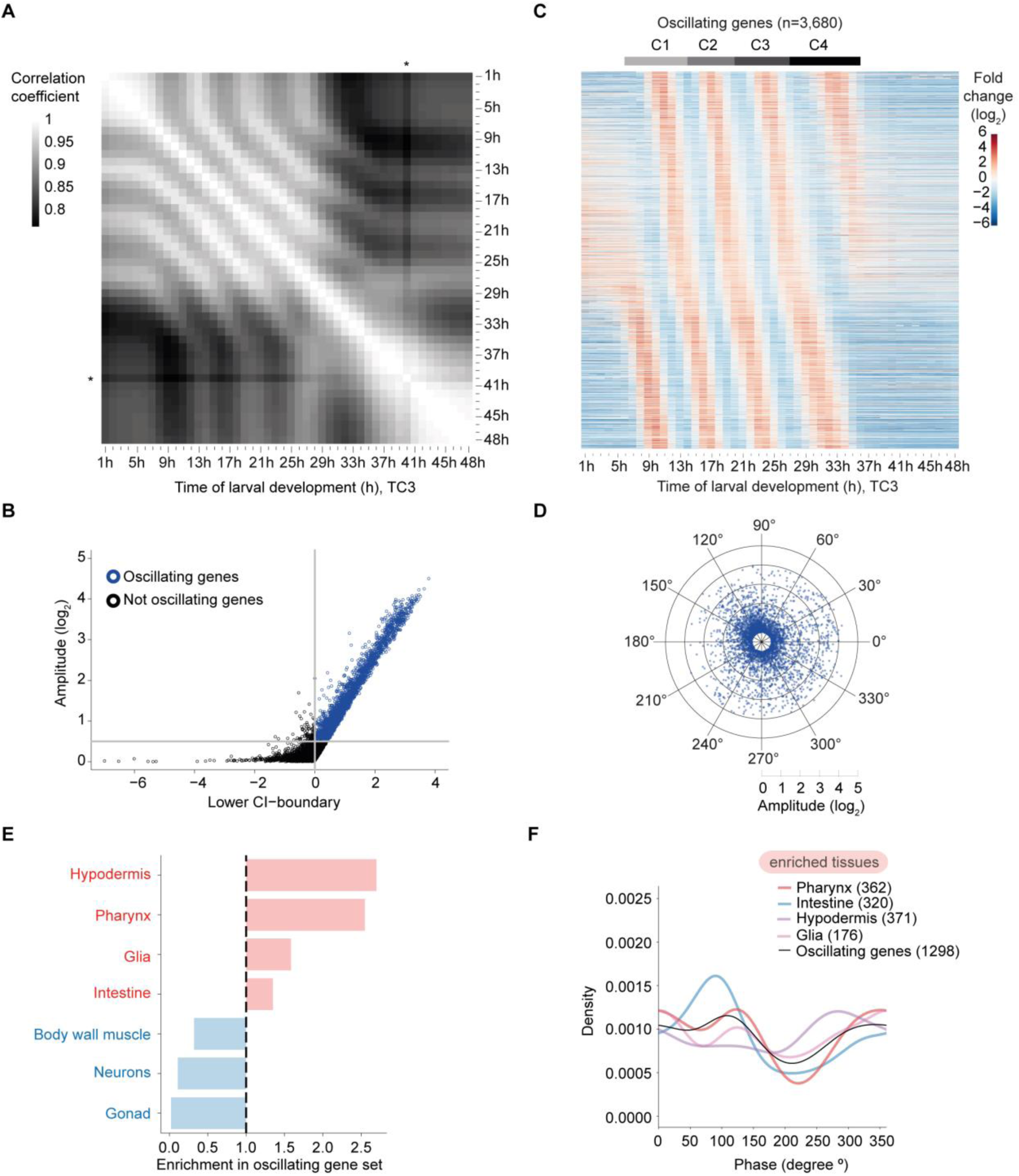
A massive oscillator with dispersed peak phases in several tissues. (A) Pairwise correlation plot of log_2_-transformed gene expression patterns obtained from synchronized population of L1 stage larvae sampled and sequenced from t = 1 h until t = 48 h (TC3; a fusion of the two time courses TC1 and TC2 after 13 h; Fig. S1A, B). Asterisk indicates an outlier, time point t = 40 h. (B) Scatter plot identifying genes with oscillatory expression (henceforth termed oscillating genes, *blue*) based on amplitude and 99% confidence interval (99%-CI) of a cosine fitting of their expression quantified in TC2 (Methods). A lower CI-boundary ≥ 0, i.e. p-value ≤ 0.01, and a log_2_(amplitude) ≥ 0.5, which corresponds to a 2-fold change from peak to trough, were used as cut-offs. Genes below either cut-off were included in the ‘not oscillating’ group (*black*). Fig. S1C reveals gene distributions in a density scatter plot. (C) Gene expression heatmaps of oscillating genes as classified in Fig. 1B and S1C. Oscillating genes were sorted by peak phase and mean expression per gene from t = 7 h to t = 36 h (when oscillations occur) was subtracted. n=3,680 as not all genes from the long time course (TC2) were detected in the early time course (TC1). Gray horizontal bars indicate the individual oscillation cycles, C1 through C4 as later determined in Fig. S8. (D) Radar chart plotting amplitude (radial axis, in log_2_) over peak phase (circular axis, in degrees) as determined by cosine fitting in Fig. 1B. (E) Enrichment (*red*) or depletion (*blue*) of tissues detected among oscillating genes expressed tissue-specifically relative to all tissue-specific genes using annotations derived from (Cao et al., 2017). Significance was tested using one-sided binomial tests which resulted in p-values < 0.001 for all tissues. (F) Density plot of the observed peak phases of tissue-specifically expressed oscillating genes for all enriched tissues.

The larger dataset enabled us to improve on the previous identification of genes with oscillatory expression (Hendriks et al., 2014). Using cosine wave fitting, and an amplitude cut-off of 2_0.5_, we classified 3,739 genes (24 % of total expressed genes) as ‘oscillating’ (i.e., rhythmically expressed) from TC2 (Fig. 1B, S1C and Table S1; Methods). Relative to the previous result of 2,718 oscillating genes (18.9% of total expressed genes) in mRNA expression data of L3 and L4 animals (Hendriks et al., 2014), this adds 1,240 new genes and excludes 219 of the previously annotated oscillating genes. We consider this latter group to be most likely false positives from the earlier analysis, resulting from the fact that some genes behave substantially different during L4 compared to the preceding stages as shown below.

Visual inspection of a gene expression heatmap of the fused time course (TC3; Fig. 1C) revealed four cycles of gene expression for the oscillating genes. Oscillations were absent during the first few hours of larval development as well as in adulthood, from ∼37 hours on, and both their onset and offset appeared to occur abruptly. We will analyze these and additional features of the system and their implications in more detail in the following sections.

### ‘Oscillating’ genes are expressed in several tissues with dispersed peak phases

An examination of the calculated peak phases confirmed the visual impression that individual transcripts peaked at a wide variety of time points, irrespective of expression amplitude (Fig. 1D). In circadian rhythms, peak phase distributions are typically clustered into three or fewer groups when examined in a specific tissue (Koike et al., 2012; Korenčič et al., 2014). However, the identity of oscillating genes differs across cell types and tissues, and for those genes that oscillate in multiple tissues, phases can differ among tissues (Zhang et al., 2014). Hence, we wondered whether the broad peak phase distribution was a consequence of our analysis of RNA from whole animals, whereas individual tissues might exhibit a more defined phase distribution.

To understand in which tissues oscillations occur, we utilized a previous annotation of tissue-specifically expressed genes (Cao et al., 2017). 1,298, and thus a substantial minority (∼35%) of oscillating genes, fell in this category for seven different tissues. They were strongly (∼2.5-fold) enriched in the hypodermis (epidermis) and pharynx, and more modestly (≤1.5-fold) in glia and intestine (Fig. 1E). By contrast, oscillating genes were greatly depleted from body wall muscle, neurons, and gonad. Hence, oscillatory gene expression occurs indeed in multiple tissues. However, although peak phase distributions deviated for each tissue to some degree from that seen for all oscillating genes, they were still widely distributed for each individual tissue (Fig. 1F).

We conclude that a wide dispersion of peak phases appears to be an inherent oscillator feature rather than the result of a convoluted output of multiple, tissue-specific oscillators with distinct phase preferences.

### Oscillations initiate with a time lag in L1

The observation that oscillations were undetectable during the first few hours of larval development and started only after > 5 hours into L1 (Fig. 1A, C) surprised us. Hence, we performed a separate experiment that covered the first 24 hours of larval development (TC4). This confirmed our initial finding of a lack of oscillations during the first few hours of larval development (Fig. 2A, B).

**Fig 2:**
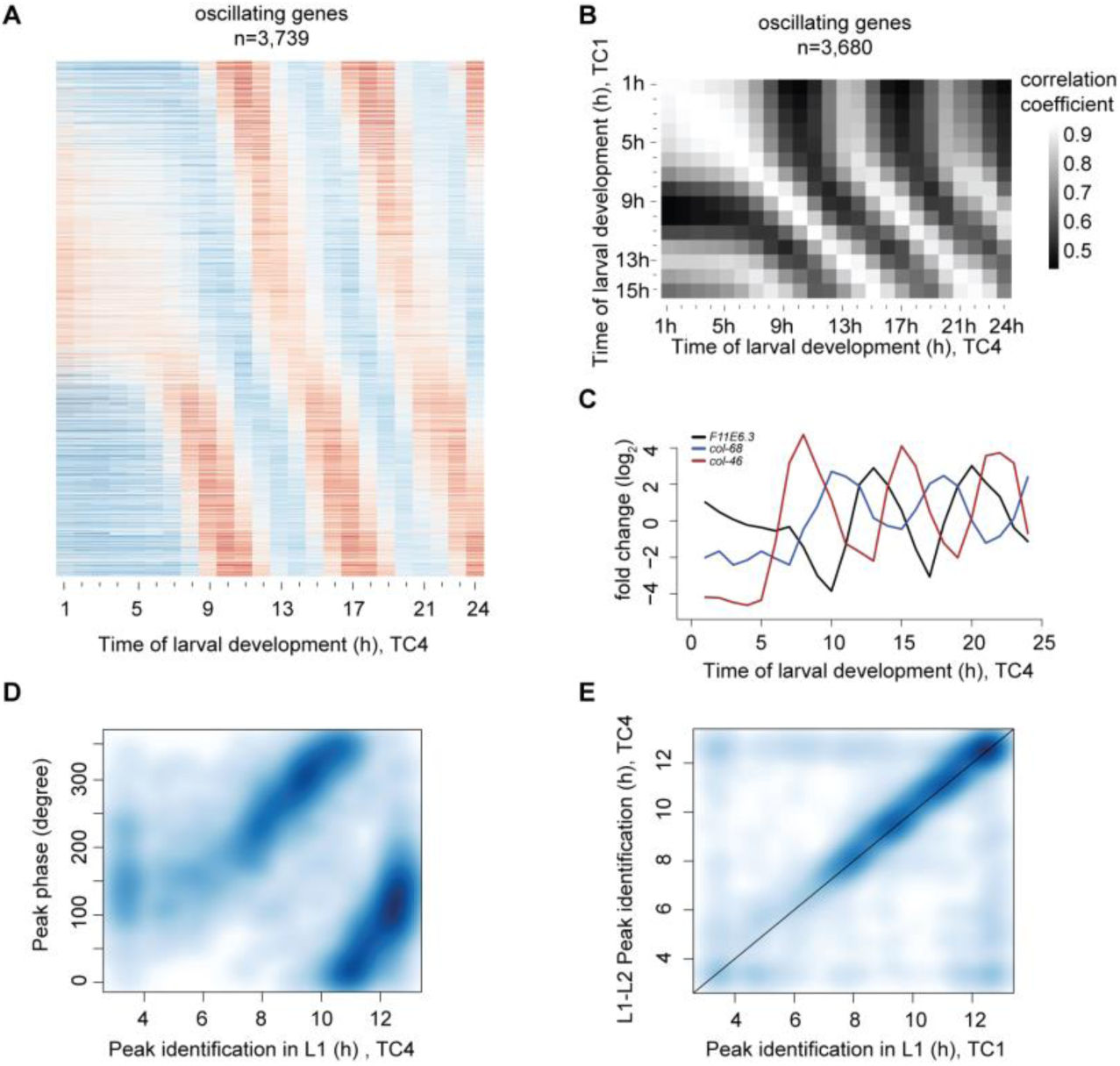
Oscillations start with a time lag in L1. (A) Gene expression heatmap of detectably expressed oscillating genes sampled from a separate early developmental time course (TC4; t = 1 h to t = 24 h). Genes were ranked according to their peak phase as determined in Fig. 1. (B) Pairwise correlation plot of log_2_-transformed oscillating gene expression data obtained from both early larval development time courses, TC1 and TC4. (C) Gene expression traces of representative genes *F11E6.3*, *col-68* and *col-46*. (D) Scatter plot of calculated oscillating gene peak phase (Fig. 1) over the time of occurrence of the first expression peak in L1 larvae, observed in TC4. Peak detection was performed using a spline analysis. As visual inspection did not reveal peaks in the heatmap during the first three hours, and as the first cycle ends at 13 h, we performed this analysis for t = 3 h to t = 13 h to reduce noise. (E) Scatterplot comparing experimentally identified first peaks of gene expression (as in D) of the two early time course replicates, TC1 and TC4. All analyses for oscillating genes identified in Fig. 1 with detectable expression (n=3,680).

To understand how oscillations initiate after the initial quiescence, we looked at individual genes and observed that the start of detectable oscillations differed for individual genes (Fig. 2A, C). Nonetheless, the occurrence of first peaks was globally well correlated to the peak phases calculated from data in Fig. 1 (Fig. 2D, E). Moreover, the transcript levels of many genes with a late-occurring (11 – 13 hours) first peak proceeded through a trough before reaching their first peak as exemplified in Fig. 2C for *F11E6.3*. We conclude that initiation of larval development after hatching is accompanied by a time lag prior to transition into an oscillatory state, and that oscillations exhibit a structure of phase-locked gene expressing patterns as soon as they become detectable.

### L1 larvae undergo an extended intermolt

Although the gene expression oscillations occur in the context of larval development, functional connections have been lacking. However, genes encoding cuticular components were reported to be enriched among previously identified oscillating genes (Hendriks et al., 2014; Kim et al., 2013), and Gene Ontology (GO-) term analysis of the new extended set of oscillating genes confirms that the top 12 enriched terms all linked to cuticle formation and molting, or protease activity (Fig. 3A). These findings, and the fact that molting is itself a rhythmic process, repeated at the end of each larval stage, suggest the possibility of a functional link between molting and gene expression oscillations.

**Fig 3:**
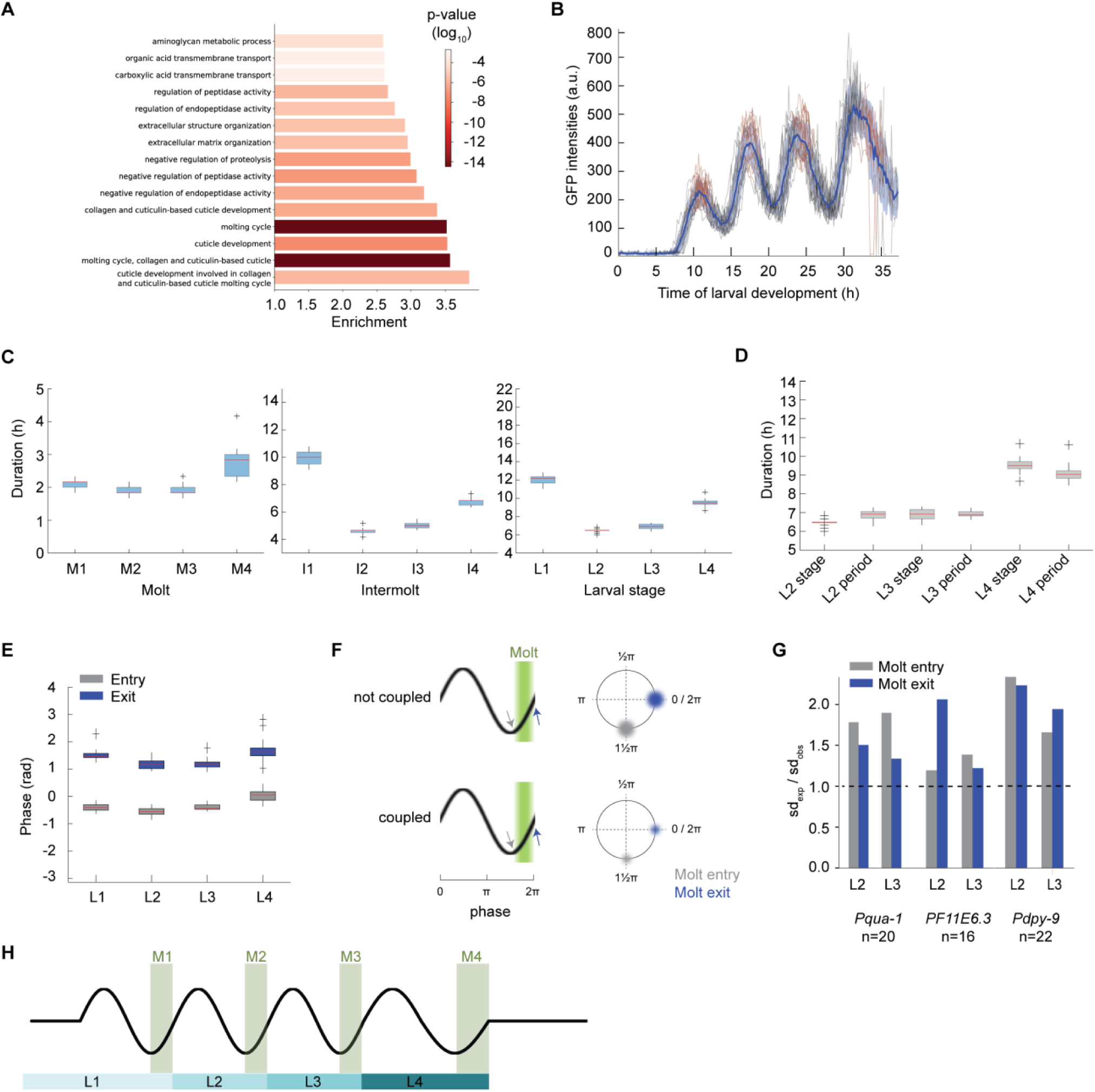
Oscillatory gene expression is coupled to molting. (A) GO-term enrichments for oscillating genes as classified in Fig. 1C. The top 15 enriched terms are displayed. (B) GFP signal quantification for *Pqua-1::gfp::pest::h2b::unc-54_3’UTR_* expressing single animals (HW2523, n=20) over larval development, starting from hatch (t = 0 h). Molts (*red*), mean intensity (*blue line*) and standard deviation across population (*shading*) are indicated. (C), (D) Boxplots of molt, intermolt and larval stage durations (C) and of larval stage durations and period times of oscillations (D) of single animals (HW2523) developing in microchambers (n=20). In (D), L1 was excluded because of the time lag before oscillations manifest after hatching. (E) Boxplot of phase at molt entry (start of lethargus) and molt exit (end of lethargus) separated by larval stages for single animals (HW2523) developing in microchambers (n=20) (F) Schematic models of size of expected phase variation (radius of colored blur) at molt entry (*gray*) and molt exit (*blue*) depending on the coupling status between oscillations and molting. (G) Barplots displaying the ratio of expected standard deviation over observed standard deviation for phase calling at either molt entry or molt exit for the indicated reporters. A dashed line indicates parity. (H) Schematic depiction of coordination between oscillatory gene expression and development. Boxplots in (C) – (E) extend from first to third quartile with a line at the median, outliers are indicated with a cross, whiskers show 1.5*IQR.

If such a link were true, we would predict that the initial period of quiescence in the early L1 stage be accompanied by a lengthened stage, and, specifically, intermolt duration. Indeed, using a luciferase-based assay that reveals the period of behavioral quiescence, or lethargus, that is associated with the molt (Fig. S2A-C), others had previously reported an extended L1 relative to other larval stages (Olmedo et al., 2015). However, they reported an extension of both molt and intermolt.

As the previously used luciferase-expressing transgenic strains developed relatively slowly and with limited synchrony across animals, presumably due to their specific genetic make-up, we repeated the experiment with a newly generated strain that expressed luciferase from a single copy integrated transgene and that developed with improved synchrony and speed (Fig. S2D-J, Methods). Our results confirmed that L1 was greatly extended relative to the other larval stages (Fig. S2I). However, in contrast to the previous findings (Olmedo et al., 2015), but consistent with our hypothesis, the differences appeared largely attributable to an extended intermolt (Fig. S2G). The duration of the first molt (M1) was instead comparable to that of M2 and M3 (Fig. S2H).

Thus, an extended first intermolt coincides with the fact that no oscillator activity can be detected by RNA sequencing during the first five hours of this larval stage. Moreover, because we performed the experiment by hatching embryos directly into food, we can conclude that the extended L1 stage is an inherent feature of *C. elegans* larval development, rather than a consequence of starvation-induced synchronization.

### Development is coupled to oscillatory gene expression

The luciferase assay revealed that also the L4 stage took significantly longer than the two preceding stages, though not as long as L1 (Fig. S2I). In this case, both the fourth intermolt and the fourth molt were extended (Fig. S2G, H). As apparent from the gene expression heatmap, and quantified below, the oscillation period during L4 was also extended. Hence, grossly similar trends appeared to occur in larval stage durations and oscillation periods, determined by the luciferase assay and RNA sequencing, respectively. We considered this as further evidence for a coupling of the two processes.

To test this hypothesis explicitly, we sought to quantify the synchrony of oscillatory gene expression and developmental progression in individual animals at the same time. To this end, we established a microchamber-based time-lapse microscopy assay by adapting a previous protocol (Turek et al., 2015). In this assay, animals are hatched and grown individually in small chambers where they can be tracked and imaged while moving freely, enabling their progression through molts. Using Mos1-mediated single copy transgene integration (MosSCI) (Frøkjær-Jensen et al., 2012), we generated transgenic animals that expressed destabilized *gfp* from the promoter of *qua-1*, a highly expressed gene with a large mRNA level amplitude.

Consistent with the RNA sequencing data, we detected oscillations of the reporter with four expression peaks (Fig. 3B). Moreover, we observed similar rates of development as in the luciferase assays when we curated the molts (Fig. 3C, Table S2, Methods). The stage durations were in good agreement with the averaged oscillation period times for each cycle, obtained through a Hilbert transform of GFP intensities, for all three larval stages, L2 through L4, for which oscillations period lengths could be reliably determined (Fig. 3D).

Single animal imaging enabled us to ask when molts occurred relative to oscillatory gene expression, and we observed a very uniform behavior across animals (Fig. 3B). To quantify this relationship, we determined the gene expression phases at molt entries and exits. We obtained highly similar values across worms within one larval stage (Fig. 3E), and only a minor drift when comparing phases across larval stages. Two additional reporter transgenes, based on the promoters of *dpy-9* and *F11E6.3*, which differ in peak expression phases from *qua-1* and one another, yielded similar results (Fig. S3).

We considered two possible interpretations of the narrow distributions of oscillation phases at molt entry and exit: first, both oscillations and development could be under independent, but precise temporal control. In this model, certain developmental events would merely coincide with specific phases of oscillations rather than being coupled to them. Therefore, variations in the periods of oscillation and development would add up, non-linearly, to the experimentally observed phase variations. Second, phase-locking of oscillatory gene expression and developmental events might result from the two processes being truly coupled and/or from one driving the other. In this case, the variations in the two periods would partially explain each other, causing a reduction in the expected phase variation relative to the first scenario (Fig. 3F).

To distinguish between these scenarios, we used error-propagation to calculate the expected error for two independent processes (Methods). Focusing on L2 and L3 stages to exclude any edge effects on period calculation by Hilbert transform, we found that this calculated error consistently exceeded the experimentally observed error (Fig. 3G), for all three reporter genes, for both molt entry and molt exit, and for both larval stages. Thus, our observations agree with the notion that development and oscillatory gene expression are functionally coupled (Fig. 3H), and potentially causally connected.

### Quantification of amplitude and period behavior over time reveal characteristic systems properties

Consistent with the coupling between oscillations and development, both the last larval stage and the period of the last oscillation cycle appeared increased (Fig. 3D), before oscillations ceased. The characteristics of such a transition from oscillatory to non-oscillatory state, or bifurcation, can be examined in the light of bifurcation theory. Bifurcation, that is, a qualitative change in system behavior, occurs in response to a change in one or more control parameters. Depending on the system’s topology, characteristic changes of amplitude and period occur during bifurcation (Fig. 4A) (Izhikevich, 2000; Salvi et al., 2016; Strogatz, 2015). Thus, transition into a quiescent state through a supercritical Hopf (supH) bifurcation involves a declining, and ultimately undetectable, amplitude and a constant period. By contrast, a Saddle Node on Invariant Cycle (SNIC) bifurcation results in a declining frequency (and thus increasing period) but a stable amplitude.

**Fig 4:**
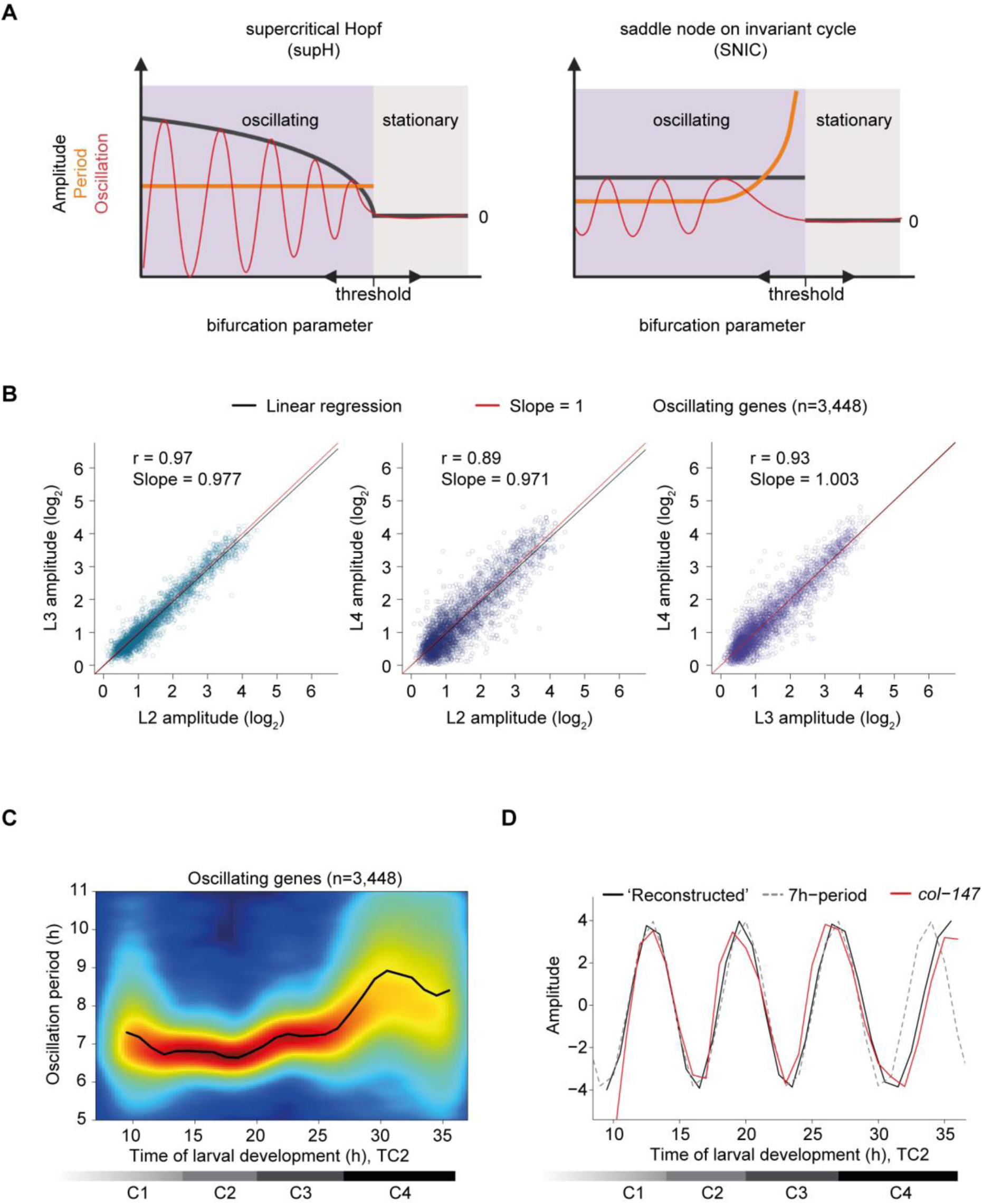
Change in period without noticeable change in amplitude. (A) Schematic depiction of amplitude and period behaviors in response to a control parameter change for an oscillatory system transitioning between a quiescent (stationary) and an oscillatory state through the indicated bifurcations (created with BioRender.com). Note that transitions can occur in either direction. (B) Amplitudes derived from cosine fitting to the individual oscillations of L2, L3 and L4 stage (TC2) plotted against each other. Pearson correlation coefficient r, slope of the linear regression (*black*) and the diagonals (slope=1; *red*) are indicated. 291 genes were excluded from oscillating genes due to altered mean expression in L4, see Fig. S4, i.e., n=3,448. (C) Density plot showing oscillation period at every time point for each of the oscillating genes (n=3,448) as quantified by Hilbert transform. Mean oscillation period over all oscillating genes is shown by the black line. Horizontal gray bars indicate oscillation cycles C1 through C4 as in Fig. 1C. (D) Expression changes for an oscillation with a constant 7h period (*dotted line*), and an oscillation reconstructed from the mean oscillation period in (C) (*black line*), both amplitudes set to four. The expression of a representative gene, *col-147* (mean normalized) is shown (*red line*). Horizontal gray bars indicate oscillation cycles C1 through C4 as in Fig. 1C.

Hence, to gain a better understanding of the oscillator’s bifurcation, we quantified oscillation amplitudes and periods over time. To minimize variations from differences between experiments, we did this for the contiguous 5 – 48 hour time course (TC2). This enabled reliable quantification of these features for the last three oscillation cycles, C2 through C4, which begin at 14 h (C2), 20 h (C3) and 27 h (C4), respectively (Fig. 4C). Excluding a small set of 291 genes that exhibited unusual expression trends during the fourth larval stage, i.e., a major change in mean expression levels (Fig. S4) this analysis revealed a good agreement of amplitudes across stages, and in particular no indication of damping during the last cycle, C4 (Fig. 4B).

We used a Hilbert transform (Pikovsky et al., 2001) to quantify the period over time with high temporal resolution, i.e., at every hour of development. The mean oscillation period thus calculated was approximately seven hours during the first cycles but increased during the fourth cycle (Fig. 4C). This change was also apparent when we reconstructed an oscillation from the mean observed oscillation period and compared it to an oscillation with a constant period of seven hours (Fig. 4D).

In summary, these analyses reveal a sudden loss of oscillation upon transition to adulthood without prior amplitude damping and an oscillator that can maintain a stable amplitude in the presence of period changes. These are features of an oscillator operating near a SNIC rather than a supH bifurcation (Fig. 4A) (Izhikevich, 2000; Salvi et al., 2016; Strogatz, 2015).

### Arrest of the oscillator in a specific phase upon transition to adulthood

SNIC and supH bifurcations differ not only in amplitude and period behavior, but also in the stable state, or fixed point, that the systems adopts. In a supH bifurcation, the system spirals from a limit cycle onto a fixed point, whereas in a SNIC bifurcation, the fixed point emerges on the limit cycle (Fig. 5A) (Saggio et al., 2017). In other words, a quiescent oscillator near a SNIC bifurcation adopts a state similar to that of a specific phase of the oscillator; the oscillator has become ‘arrested’. By contrast, following a supH bifurcation, the oscillator adopts a stable state that is distinct from any phase of the oscillator. Hence, if the *C. elegans* oscillator entered an arrested state through a SNIC bifurcation, the overall expression profile of the oscillating genes in the adult stage should resemble that seen at some other time point during larval development.

**Fig 5:**
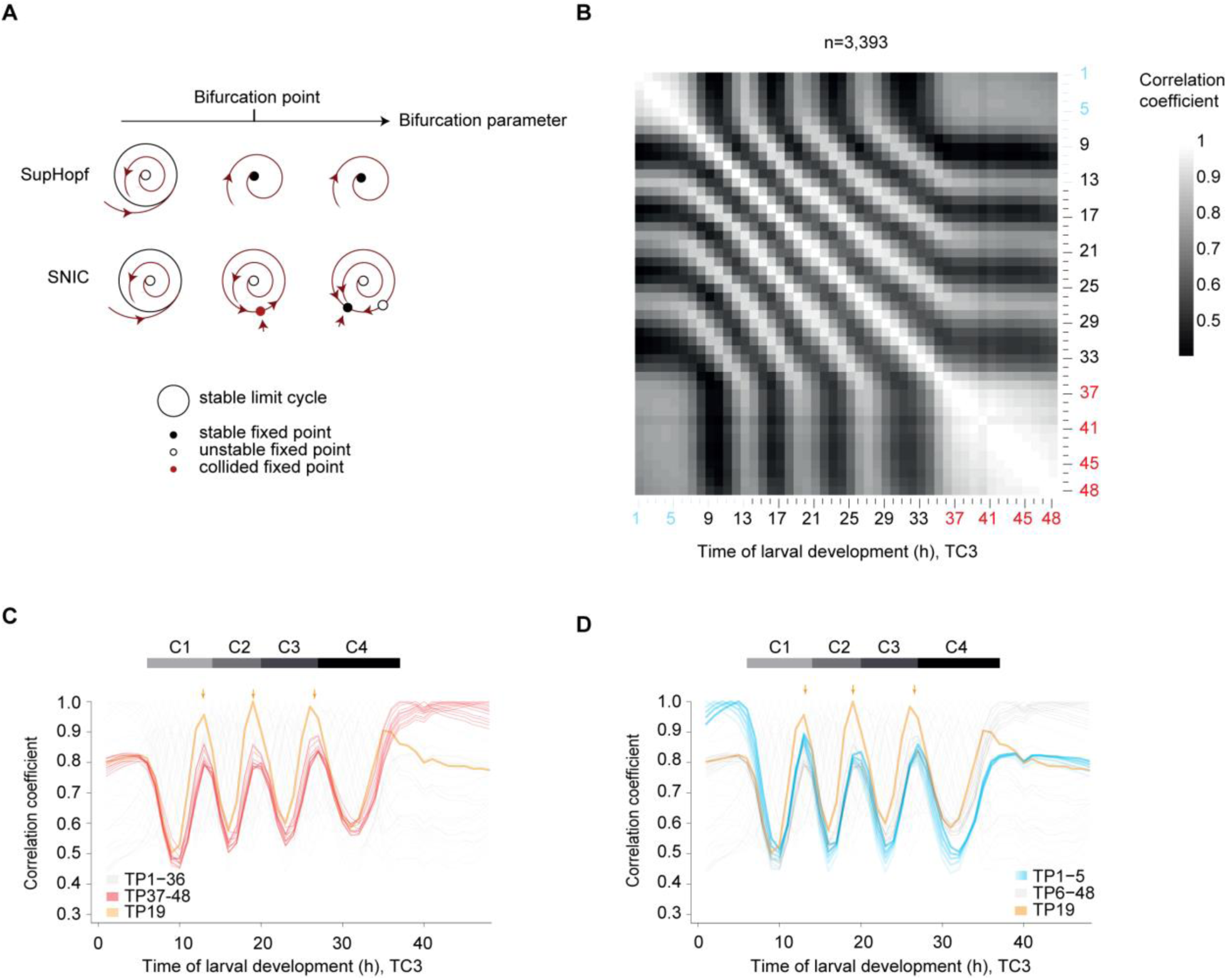
The oscillator is phase arrested in early L1 and adults. (A) Phase plane diagrams depicting supH and SNIC bifurcations, respectively, showing the change in qualitative behavior as the bifurcation parameter value changes (arrow). The bifurcation point, i.e. the parameter value at which the bifurcation occurs, is indicated. (B) Pairwise correlation plot of log_2_-transformed oscillating gene expression data obtained from TC3, i.e., the fusion of TC1 (*blue labels*) and TC2 (*black*). Genes which deviated in mean expression in L4 were excluded (Fig. S4), resulting in n=3,393 genes. (C) Lines of correlation for TP37–48 (*red*) to all time points in the fused larval time course. Arrows indicate local correlation maxima at TP13, 19 and 26/27. Correlation traces for TP ≤ 36 h are shown in light gray, except for TP19 (*orange*). Fig. S5 illustrates how correlation lines were generated. (D) Lines of correlation for TP1–5 (*blue*) and TP19 (*orange*) to all time points in the fused larval time course. Arrows indicate local correlation maxima at TP13, 19 and 26/27. All correlations were determined by Pearson correlation. Correlation values were obtained from Fig. 5B.

To test this prediction, we analyzed the correlation of oscillating gene expression for adult time points (TP ≥ 37 h) to all other time points of the fused time course (TC3). (In the following, we will use “TPx” to refer to any time <u>p</u>oint ‘x’, in hours, after hatching. Technically, this is defined in our experiment as the time after plating synchronized, first larval stage animals on food.) For this analysis (illustrated in Fig. S5), we used the pairwise correlation matrix resulting from the oscillating gene set without the previously excluded genes that changed in expression in the L4 stage (Fig 5B). This provided two insights. First, correlation coefficients among adult time points all exceeded 0.8 with little change over time, confirming the high similarity of samples TP37 – 48 to one another and thus an absence of detectable oscillations. Second, in addition to one another, TP37 – 48 are particularly highly correlated to a specific time – and thus phase – of the oscillatory regime, namely TP13 and the ‘repetitive’ TP19 and TP26/27 (Fig. 5C, arrows). In other words, expression levels of oscillating genes in the adult resembled a specific larval oscillator phase, providing further support for a SNIC bifurcation.

### Phase-specific arrest of the oscillator after hatching

We noticed that the gene expression states of TP37 – 48 also correlated well to each of TP1 – 5; i.e., the early L1 larval stage (Fig. 5B). To examine this further, we performed the same correlation analysis as described above, but now for TP1 – 5. Mirroring the adult situation, correlation coefficients among all these five time points were high and exhibited little change over time, and TP1 – 5 exhibited particularly high levels of correlation to TP13 and the ‘repetitive’ TP19 and TP27 (Fig. 5D). These are the same larval time points to which the adult time points exhibit maximum similarity. We confirmed these two key observations when fusing the independent time course TC4 to TC2 (generating TC5; Fig. S6).

We conclude that also during the first five hours after plating, oscillating genes adopt a stable expression profile that resembles a specific phase of the oscillator. In other words, both the transition into the oscillatory state during L1 and out of it during L4 occur through a SNIC bifurcation. This finding indeed explains our observation (Fig. 2) that in L1 stage larvae, oscillations exhibit a structure of phase-locked gene expressing patterns as soon as they become detectable: the oscillator initiates from an arrested phase.

### Initiation of oscillation soon after gastrulation

We wondered how the oscillator entered the arrested state observed in early larvae, i.e., what dynamics the class of larval oscillating genes exhibited in embryos. Hence, we examined single embryo gene expression data from a published time series (Hashimshony et al., 2015). When plotting the embryonic expression patterns of oscillating genes sorted by their peak phase defined in larvae, we observed a dynamic expression pattern with a striking phase signature (Fig. 6A). To investigate this further, we performed a correlation analysis between embryonic and larval time points (TC3) for the oscillating genes (Fig. S7A). When we plotted the correlation coefficients for each embryonic time point over larval time we observed two distinct behaviors (Fig. 6B, C), which separated at ∼380 min (95%-CI: 317.6 minutes – 444.2 minutes) (Fig. 6D; Fig. S7B): First, from the start of embryogenesis until ∼380 min, the peak of correlation occurred always for the same larval time point, but the extent of correlation increased rapidly (Fig. 6B, D). Second, past ∼380 min of embryonic development, the peaks of correlation moved progressively from TP14 (which we define as 0°/360° because it demarcates the end of the first and the beginning of the second oscillation cycle in the fused time course, Fig. 1C); towards TP19 (accordingly defined as 300°), but the extent of correlation increased only modestly (Fig 6C, D).

**Fig 6:**
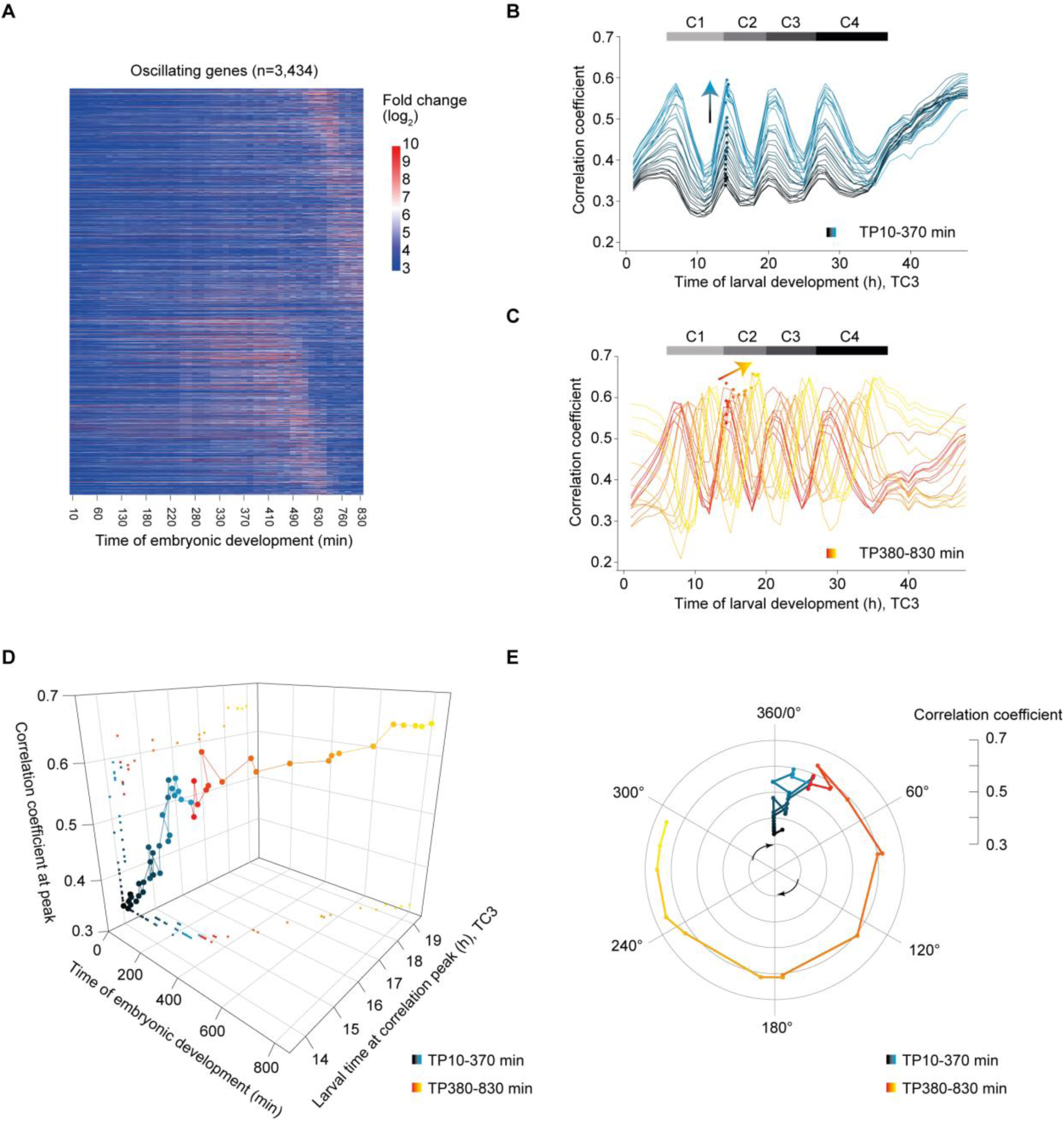
Transition to an oscillatory state during embryogenesis. (A) Heatmap of log_2_-transformed embryonic expression of oscillating genes, excluding L4 deviating genes, sorted by larval peak phase (defined as in Fig. 1). (B, C) Pairwise correlation coefficients between embryonic and larval time points (Fig. S7) plotted over larval time for embryonic TP10-370 min (B, *black-blue gradient*) and TP380-830 min (C, *red-yellow gradient*), respectively. Dots represent peaks of the correlation lines after spline analysis in the second oscillation cycle (C2), arrows indicate trends. Horizontal gray bars indicate oscillation cycles C1 through C4 as in Fig. 1C. (D) 3D-scatter plot of the correlation coefficient peak for each embryonic time point to the time of larval development at the second oscillation cycle (C2). Embryonic time is determined by time of sample collection, larval time by spline interpolation. Color scheme as in B and C. (E) Polar plot of correlation coefficient peak over the time point in the second larval oscillation cycle (C2) at which the correlation peak is detected. TP14 is defined as 0° and correlates most highly to TP20, thus defined as 360°. Values are as in D; Color scheme as in B and C. All correlations were determined by Pearson correlation.

We conclude that the system adopts two distinct states during embryogenesis (Fig. 6E): Initially, it approaches the oscillatory regime through increasing similarity to the oscillator phase TP14/0°. After completion of gastrulation and around the beginning of morphogenesis/organogenesis (Hall et al., 2017), it transitions into the oscillatory state and reaches, at hatching, a phase corresponding to larval ∼TP19/300°, where oscillations arrest until resumption later in L1.

### A shared oscillator phase for experimentally induced and naturally occurring bifurcations

The arrested states of the oscillator in both early L1 stage larvae and in adults are highly similar and resemble the oscillator state at TP19/300°. Therefore, we wondered whether this oscillator phase was particularly conducive to state transitions of the system in response to changes in the developmental trajectory. To test this, we examined animals that exited from dauer arrest, a diapause stage that animals enter during larval development under conditions of environmental stress such as heat, crowding, or food shortage. Using a published time course of animals released from dauer arrest after starvation (Hendriks et al., 2014), we found that their expression patterns of oscillating genes correlated highly with those of animals initiating oscillations (TC3) in the L1 stage (Fig. 7A, B). Additionally, gene expression patterns at 1 hour through 5 hours and at 13 hours post-dauer were highly correlated to those of the repetitive TP13, TP19 and TP26/27 during continuous development. Hence, the system state during a period of quiescence during the first five hours after placing animals on food corresponds to an arrest of the oscillation in a phase seen at TP19/300° of the continuous development time course.

**Fig 7:**
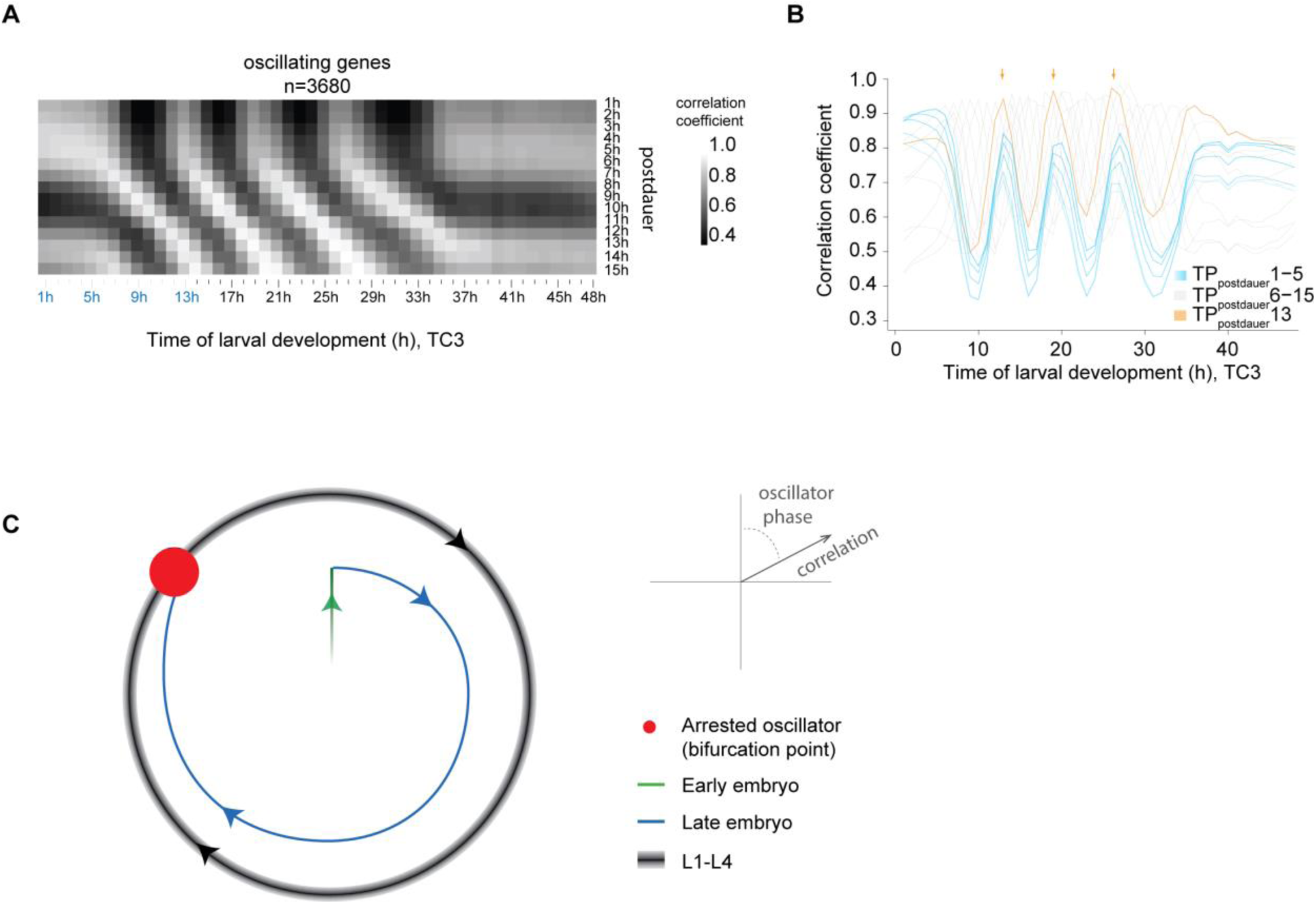
Re-initiation of oscillations after dauer from an arrested oscillator phase. (A) Pairwise-correlation map of log_2_-transformed oscillatory gene expression of dauer exit samples (“postdauer”) and fused larval time course (TC3) samples. (B) Correlation of the indicated time points after plating dauer-arrested animals on food (TPpostdauer) to the fused larval time course, TC3. Arrows indicate peaks of correlation to TP13/19/26.5 (300°) of TC3. (C) Schematic depiction of behavior of the *C. elegans* oscillator from embryo to adult. A phase-specific arrest (*red dot*) is observed in hatch, early L1, young adults, and dauer-arrested animals. See Fig. S8 for additional data supporting four similar oscillation cycles during L1 through L4.

We conclude that the system bifurcates in the same manner during continuous, unperturbed development, after hatching, and in response to a perturbation, namely starvation-induced dauer arrest.

## Discussion

In this study, we have characterized biological function and behavior of the *C. elegans* larval oscillator. Our results from single animal- and population-based analyses reveal a close coupling to development, and specifically molting, and imply that processes essential for molting may not be restricted to lethargus. We have observed that oscillations are highly similar during the four cycles (Fig. 7C, Fig. S8). Yet, oscillations cease and (re-) initiate several times during physiological development, and similar state transitions, or bifurcations, of the system can be induced through an external perturbation (Fig. 7C). In particular, all non-oscillatory states correspond to an arrest of the oscillator in one specific phase. Hence, the observed bifurcations provide a conceptual model of how a developmental checkpoint can operate to halt larval development at a particular, repetitive point of development. Moreover, they constrain possible system architectures and properties as well as the choice and parametrization of mathematical models that can represent the system.

### Oscillatory expression of thousands of genes

Our previous work (Hendriks et al., 2014) identified ∼2,700 oscillating (i.e., rhythmically expressing) genes, a number that we now increase to 3,739 genes (24% of total expressed genes). We attribute this improved identification of oscillating genes to the fact that our present analysis focused on the L1, L2 and L3 stages, where a constant oscillation period of ∼7 hours facilitates cosine wave fitting. This contrasts with the situation in the previous experiment, which used data from the L3 and L4 stages and thus, as we reveal here, a time of changing period.

Even our current estimate is conservative, i.e., the ‘non-oscillating’ genes contain genes that exhibit oscillatory expression with low amplitude or, potentially, strongly non-sinusoidal shapes. It is possible that such dynamics may play important roles for specific genes and processes and our data provide a resource to identify these in the future. However, here we focused on genes with robust and extensive oscillations to facilitate functional dissection of the oscillator.

### A developmental oscillator with functions in and beyond molting and lethargus

The physiological function of the *C. elegans* oscillator has remained unclear. Here, we have tested a possible connection with molting. By quantifying the periods of both molting and gene expression oscillations simultaneously, in the same individual animals, we revealed their high degree of similarity and showed that the two processes are more closely phase-locked than expected from mere coincidence, i.e., they are coupled. We propose that a function of the oscillator as a developmental clock provides a parsimonious explanation for the coupling, but other models remain possible, e.g., the oscillator may facilitate an efficient molting process by anticipating the time of peak demand for cuticular building blocks and other factors.

Conventionally, molting is subdivided into three distinct steps, namely apolysis (severing of connections between the cuticle and the underlying epidermis), new cuticle synthesis, and ecdysis (cuticle shedding) (Lažetić and Fay, 2017). The first two occur during, and the latter terminates, lethargus, a period of behavioral quiescence. However, we find that the temporal structure imposed by the *C. elegans* oscillator extends beyond lethargus. Our data reveal two probable causes: occurrence of processes required for molting before lethargus and a temporal organization that extends to processes unrelated to molting.

Specifically, we observed initiation of oscillations in embryos, which execute cuticle synthesis but neither apolysis nor ecdysis, at ∼380 min into embryo development and thus long before the first signs of cuticle synthesis at ∼600 min (Sulston et al., 1983). Instead, this time coincides with formation of an apical extracellular matrix (ECM). Although termed embryonic sheath, we find genes encoding components of this ECM, namely *sym-1; fbn-1; noah-1; noah-2* (Vuong-Brender et al., 2017), also detectably expressed in larvae, and all four are required for larval molting or proper cuticle formation (Frand et al., 2005; Niwa et al., 2009). Moreover, their mRNA levels oscillate with high amplitudes, and as predicted from the embryonic data, their expression peaks long before lethargus, and in fact shortly after ecdysis, i.e., <u>after</u> a molt has been completed. Hence, to account for all these facts, we propose that molting involve processes that are executed long before the onset of lethargus and that include ECM remodeling.

However, even a substantially more complex molting process may fail to account for the fact that a majority of oscillating genes is phase-locked with the molt but exhibits peak expression outside the molt and lacks any obvious link to molting. We consider it plausible that robust larval development may benefit from a coordination of molting with other physiological or developmental processes, as previously postulated for skin cell proliferation (Ruaud and Bessereau, 2006). Similarly, as lethargus involves cessation of food-uptake, oscillatory gene expression may serve to anticipate this event, consistent with a large number of intestinally expressed oscillating genes. Nonetheless, even for this class, a broad dispersion of peak expression phases may suggest additional functions, yet to be uncovered. Whatever the benefit, it is evident that the oscillator imposes a temporal structure of gene expression that extends far beyond lethargus.

### Oscillatory state transitions and developmental checkpoints

We have observed a loss of oscillations under three distinct conditions, in early L1 stage larvae, dauer arrested animals, and adults. The similarity of the oscillator states under all three conditions is striking and involves an arrest in the same specific phase.

Formally, for the L1 arrest, we cannot distinguish between perturbation-induced or naturally occurring arrest, as the sequencing experiments required animal synchronization by hatching animals in the absence of food, causing a transient arrest of development. However, the fact that the L1 stage is extended also in animals hatched into food suggests that they may adopt a similar arrested state even in the presence of food, perhaps because the nutritional resources in the egg (i.e., egg yolk) have become depleted by the time that hatching occurs. In other words, synchronization of L1 animals by hatching them in the absence of food may propagate a pre-existing transient developmental and oscillator arrest.

Irrespective of this interpretation, a key feature of the arrests that we observe under different conditions is that they always occur in the same phase. This is a behavior one would predict for a repetitive developmental checkpoint. Such a checkpoint has indeed been found to operate shortly after each larval molt exit, arresting development in response to a lack of food (Schindler et al., 2014). Importantly, developmental arrest does not result from an acute shortage of resources. Rather, it is a genetically encoded, presumably adaptive, response to nutritionally poor conditions, as demonstrated by the fact that mutations in the *daf-2*/IGFR signaling pathway causes animals to continue development under the same food-depleted conditions (Baugh, 2013; Schindler et al., 2014).

Within the limits of our resolution, the phase of the arrested oscillator corresponds to the phase seen around ecdysis. Hence, oscillations and development are synchronously arrested, and we propose that signals related to food sensing, metabolism, or nutritional state of the animal help to control the state of the oscillatory system and thereby developmental progression. An oscillator operating near a SNIC bifurcation appears ideally suited to processing such information, because it acts as a signal integrator, i.e., it becomes active when a signal threshold is surpassed (Forger, 2017; Izhikevich, 2000). This contrasts with the behavior of oscillators operating near a supH bifurcation, which function as resonators, i.e., they respond most strongly to an incoming signal of a preferred frequency. Hence, both the phase-specific arrest and the integrator function as characteristics of an oscillator operating in the vicinity of a SNIC bifurcation are physiologically relevant features of this *C. elegans* oscillator.

### Insights into oscillator architecture and constraints for mathematical modelling

What mechanisms determine the type and behavior of the *C. elegans* oscillator? Although the nature and wiring of the ‘core oscillator’, i.e., the machinery that drives the pervasive gene expression oscillation, remains to be established, the behavior of the oscillator that we characterized here provides clear constraints. Thus, a change in period without a noticeable change in amplitude, as seen in L4 stage larvae, is a feature of rigid oscillators (Abraham et al., 2010) that is considered incompatible with the function of a simple negative feedback loop but compatible with the operation of interlinked positive and negative feedback loops (Mönke et al., 2017; Tsai et al., 2008). This conclusion is supported by evidence from synthetic biology, where most synthetic genetic oscillators appear to operate near a supH bifurcation (Purcell et al., 2010). An exception are so-called amplified negative feedback oscillators, which operate near a SNIC bifurcation and rely on interlinked negative and positive feedback loops.

Beyond constraining possible oscillator architectures, our experimental observations will help to guide mathematical modelling of the *C. elegans* oscillator. Modelling is needed because the nonlinear dynamic behaviors of oscillators are difficult to grasp intuitively. However, it is usually difficult to ensure the relevance of a given model, because both its formulation and its parametrization determine whether oscillations occur and which behaviors the resulting oscillator model displays. Amplified negative feedback oscillators are a case in point as they can also operate near a supH bifurcation; operation near a SNIC bifurcation occurs only in a certain parameter space (Conrad et al., 2008; Guantes and Poyatos, 2006). The experimental characterization of the system’s bifurcation that we have achieved here will therefore provide crucial constraints to exclude invalid mathematical models.

We do note that mathematical models of somitogenesis clocks, inspired by mechanistic knowledge about the identity of individual oscillator components and their wiring, tend to represent oscillators operating near a supH bifurcation (Jensen et al., 2010; Webb et al., 2016). This appears consistent with observations on isolated cells *in vitro* (Webb et al., 2016). At the same time it contrasts with changes in both amplitude and period that were shown to occur in zebrafish embryos during somite formation and prior to cessation of oscillation (Shih et al., 2015). Thus, and because an analysis of bifurcation behavior of somitogenesis clocks *in vivo* is challenging due to a complex space-dependence of oscillation features (Soroldoni et al., 2014), it remains to be answered whether and to what extent the *C. elegans* oscillator and the somitogenesis clocks share specific properties. However, whatever the answer, a comparison of the similarities and differences in behaviors, architectures and topologies will help to reveal whether and to what extent diverse developmental oscillators follow common design principles.

## Acknowledgements

We thank Stephane Thiry, Kirsten Jacobeit and the FMI Functional Genomics Facility for RNA sequencing, Iskra Katic for help in generating transgenic strains, Maria Olmédo and Henrik Bringmann for introducing us to the luciferase and the single animal imaging assays, respectively, Dimos Gaidatzis and Michael Stadler for advice on computational analyses, Laurent Gelman for help with imaging, and Benjamin Towbin, Lucas Morales Moya, Prisca Liberali and Luca Giorgetti for comments on the manuscript.

## Funding

M.W.M.M. is a recipient of a Boehringer Ingelheim Fonds PhD fellowship. This work is part of a project that has received funding from the European Research Council (ERC) under the European Union’s Horizon 2020 research and innovation programme (Grant agreement No. 741269, to H.G.). The FMI is core-funded by the Novartis Research Foundation.

## Author contributions

Gert-Jan Hendriks and Yannick Hauser performed RNA sequencing time courses. Milou Meeuse and Yannick Hauser analyzed RNA sequencing data. Milou Meeuse performed and analyzed luciferase assays. Guy Bogaarts developed the graphical user interface for the luciferase data. Yannick Hauser acquired and analyzed single worm imaging data. Jan Eglinger wrote the KNIME workflow for the single worm imaging. Charisios Tsiairis conceived parts of the analysis. Helge Großhans, Milou Meeuse and Yannick Hauser conceived the project and wrote the manuscript.

## Competing interests

The authors declare no competing interests.

## Data and materials availability

All sequencing data generated for this study have been deposited in NCBI’s Gene Expression Omnibus (Edgar et al., 2002) and are accessible through GEO SuperSeries accession number GSE133576 https://www.ncbi.nlm.nih.gov/geo/query/acc.cgi?acc=GSE133576. A reviewer token for data access is available through the editorial office. The dauer exit time course was published previously (Hendriks et al., 2014) and is accessible through GEO Series accession number GSE52910 (http://www.ncbi.nlm.nih.gov/geo/query/acc.cgi?acc=GSE52910). Published research reagents from the FMI are shared with the academic community under a Material Transfer Agreement (MTA) having terms and conditions corresponding to those of the UBMTA (Uniform Biological Material Transfer Agreement).

## Conflict of Interests

The authors declare that they have no conflict of interest.

## Methods

### *C. elegans* strains

The Bristol N2 strain was used as wild type. The following transgenic strains were used: HW1370: *EG6699; xeSi136 [F11E6.3p::gfp::h2b::pest::unc-54 3’UTR; unc-119 +] II* (this study) HW1939: *EG6699, xeSi296 [Peft-3::luc::gfp::unc-54 3’UTR, unc-119(+)] II* (this study) HW2523: *EG6699*: *xeSi437 [Pqua-1::gfp::h2b::pest::unc-54 3’UTR; unc-119 +] II* (this study) HW2526: *EG6699: xeSi440 [Pdpy-9::gfp::h2b::pest::unc-54 3’UTR; unc-119 +] II* (this study) PE254: *feIs5 [Psur-5::luc::gfp; rol-6(su1006)] V* (Lagido et al., 2008) PE255: *feIs5 [Psur-5::luc::gfp; rol-6(su1006)] X* (Lagido et al., 2008)

All transcriptional reporters and luciferase constructs produced for this study were generated using Gibson assembly (Gibson et al., 2009) and the destination vector pCFJ150 (Frøkjaer-Jensen et al., 2008). First a starting plasmid was generated by combining *Not*I digested pCFJ150, with either *Nhe-1::GFP-Pest-H2B* or *Nhe-1::luciferase::GFP* (adapted from pSLGCV (Lagido et al., 2008)) and ordered as codon optimized, intron containing gBlocks® Gene Fragment (Integrated DNA Technologies), and *unc-54 3’UTR* (amplified from genomic DNA) to yield pYPH0.14 and pMM001 respectively. Second, promoters consisting of either 2kb upstream of the ATG or up to the next gene were amplified from *C. elegans* genomic DNA before inserting them into *Nhe*I-digested pYPH0.14 or pMM001. PCR primers and resulting plasmids are listed in the Table S3. Third, we obtained transgenic worms by single-copy integration into EG8079 worms, containing the universal *ttTi5605* locus on chromosome II by following the published protocol for injection with low DNA concentration (Frøkjær-Jensen et al., 2012). All MosSCI strains were backcrossed at least twice.

### Methods luciferase assay

Gravid adults were bleached and single embryos were transferred by pipetting into a well of a white, flat-bottom, 384-well plate (Berthold Technologies, 32505). Embryos hatched and developed in 90 µL volume containing *E. coli* OP50 (OD_600_ = 0.9) diluted in S-Basal medium (Stiernagle, Theresa, n.d.), and 100 μM Firefly D-Luciferin (p.j.k., 102111). Plates were sealed with Breathe Easier sealing membrane (Diversified Biotech, BERM-2000). Luminescence was measured using a Luminometer (Berthold Technologies, Centro XS3 LB 960) for 0.5 seconds every 10 minutes for 72 hours at 20°C in a temperature-controlled incubator and is given in arbitrary units.

Luminescence data was analyzed using an automated algorithm for molt detection on trend-corrected data as described previously (Olmedo et al., 2015), but implemented in MATLAB, and with the option to manually annotate molts in a Graphical User Interface. The hatch was identified as the first data point (starting from time point 4 to avoid edge effects) that exceeds the following value: the mean + 5*stdev of the raw luminescence of the first 20 time points.

To quantify the duration of the molts, we subtracted the time point at molt entry from the time point at molt exit. To quantify the duration of larval stages, we subtracted the time point at molt exit of the previous stage (or time point at hatch for L1) from the time point at molt exit of the current stage. The duration of the intermolt was quantified as duration of the molt subtracted from duration of the larval stage. For statistical analysis, we assumed the durations to be normally distributed and used Welch two-sample and two-sided t-test, i.e. the function ‘t.test’ of the package ‘stats’ (version 3.5.1) (R Core Team, n.d.) in R.

### RNA sequencing

For RNA sequencing, synchronized L1 worms, obtained by hatching eggs in the absence of food, were cultured at 25°C and collected hourly from 1 hour until 15 hours of larval development, or 5 hours until 48 hours of larval development, for L1–L2 time course (TC1) and L1–YA time course (TC2) respectively. A replicate experiment was performed at room temperature from 1 hours until 24 hours (TC4). RNA was extracted in Tri Reagent and DNase-treated as described previously (Hendriks et al., 2014). For the TC2 and TC4, libraries were prepared using the TruSeq Illumina mRNA-seq (stranded – high input), followed by the Hiseq50 Cycle Single-end reads protocol on HiSeq2500. For the TC1, libraries were prepared using the Illumina TruSeq mRNA-Seq Sample Prep Kit (Strand-sequenced: any), followed by the Hiseq50 Cycle Single-end reads protocol on HiSeq2500.

### Processing of RNA-seq data

RNA-seq data were mapped to the *C. elegans* genome using the qAlign function (splicedAlignment=TRUE) from the QuasR package (Au et al., 2010; Gaidatzis et al., 2015) in R. Gene expression was quantified using qCount function from the QuasR package in R. For TC2 and Dauer exit (Hendriks et al., 2014) time course, QuasR version 1.8.4 was used, and data was aligned to the ce10 genome using Rbowtie aligner version 1.8.0. For TC1, QuasR version 1.2.2 was used, and data was aligned to the ce6 genome using Rbowtie aligner version 1.2.0. For TC4 (Fig. 2), RNA-seq data were mapped to the *C. elegans* ce10 genome using STAR with default parameters (version 2.7.0f) and reads were counted using htseq-count (version = 0.11.2). Counts were scaled by total mapped library size for each sample. A pseudocount of 8 was added and counts were log_2_-transformed. For the TC2, lowly expressed genes were excluded (maximum log_2_-transformed gene expression - (log_2_(gene width)-mean(log_2_(gene width))) ≤ 6). This step was omitted in the early time courses because many genes start robust expressing only after 5-6 hours. Expression data of the dauer exit time course was obtained from (Hendriks et al., 2014).

### Classification of genes

To classify genes, we applied cosine fitting to the log_2_-transformed gene expression levels from t=10 hours until t=25 hours of developmental time (mid L1 until late L3) of TC2, when the oscillation period is most stable (Fig. 4C). During this time the oscillation period is approximately 7 hours, which we used as fixed period for the cosine fitting. We built a linear model as described (Hendriks et al., 2014) using cos(ωt) and –sin(ωt) as regressors (with 13 degrees of freedom). In short, a cosine curve can be represented as:

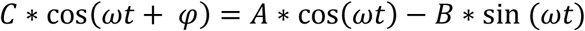

With *A* = *C* * cos(*φ*)

and *B* = *C* * sin(*φ*)

From the linear regression (‘lm’ function of the package ‘stats’ in R) we obtained the coefficients A and B, and their standard errors. A and B represent the phase and the amplitude of the oscillation:

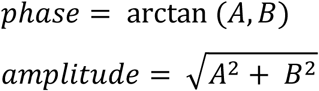

As the density of the genes strongly decreased around 0.5 (Fig. S1C) we used amplitude ≥ 0.5 as a first classifier. We propagated the standard error of the coefficients A and B to the amplitude using Taylor expansion in the ‘propagate’ function (expr=expression(sqrt(((A^2)+(B^2))), ntype = ‘stat’, do.sim=FALSE, alpha=0.01) from the package ‘propagate’ (version 1.0-6) (Spiess, 2018) in R. We obtained a 99% confidence interval (99%-CI) for each gene. As 99%-CI that does not include 0 is significant (p-value=0.01), we used the lower boundary (0.5%) of the CI as a second classifier. Thus, we classified genes with an amplitude ≥ 0.5 and lower CI-boundary ≥ 0 as ‘oscillating’ and genes with an amplitude < 0.5 or a lower CI-boundary < 0 were classified as ‘not oscillating’ (Fig. 1B, S1C). Every gene thus has an amplitude and a peak phase. A peak phase of 0° is arbitrarily chosen, and thus current peak phases are expected to differ systematically from the previously assigned peak phases (Hendriks et al., 2014). To compare the peak phases of TC2 with those of the previously published L3-YA time course (TC6), we calculated the phase difference (TC2 – TC6) (Fig. S1D, E). We added 360° to the difference and used the modulus operator (%%360), to maintain the circularity within the data. The coefficient of determination, R_2_, was calculated by 1-(SSres/SStot), in which the SStot (total sum of squares) is the sum of squares in peak phase of the L1-YA time course. SSres (response sum of squares) is the sum of squares of the phase difference.

### Time course fusion

In order to obtain an RNAseq time course spanning the complete larval development, we fused the L1–L2 time course (TC1, TP1 – TP15) with the L1–YA course (TC2, TP5 – TP48). To decide which time points to choose from the individual time courses, we correlated the gene expression of all genes (n = 19,934) of both time courses against each other using the log_2_ transformed data with a pseudocount of 8 with pearson correlation. In general, we saw good correlation between the two time courses, e.g. TP1_(TC1, L1–L2)_ correlated well with TP1_(TC2, L1–YA)_ etc. (Fig. S1B). Additionally, we could see comparable correlation values of TP13_(TC2, L1–YA)_ and TP13_(TC1, L1-L2)_ with TP1–5_(TC1, L1–L2)_ (not shown). We concluded that TP13_(TC1, L1–L2)_ and TP13_(TC2, L1–YA)_ are highly similar and thus fused at this time point, i.e., combined TP1–TP13_(TC1, L1–L2)_ with TP14–TP48_(TC2, L1–YA)_.

### Exclusion of genes based on L4 mean expression

Given that oscillating genes were identified based on gene expression in TP10-TP25, when oscillation period is most stable, some genes showed deviating behavior in the last oscillation cycle, C4. Hence, for quantification of oscillation amplitude, period and correlation, we excluded those genes. We determined the mean expression levels for each gene over time in oscillation cycles C2 (TP14-TP20), C3 (TP20-TP27) and C4 (TP27-TP36). Genes (n=291) were excluded if the absolute value of the difference in mean expression between L2 and L4 normalized by their mean difference exceeded 0.25, i.e. abs((L2meanExpr-L4meanExpr)/(0.5*(L2meanExpr-L4meanExpr)))>0.25.

### Quantification of oscillation amplitude

To quantify the oscillation amplitude for each larval stage, we split the TC2 in 4 separate cycles, roughly corresponding to the developmental stages, i.e. C1: TP6-TP14, C2: TP14-TP20, C3: TP20-TP27 and C4: TP27-TP36 developmental time. We applied cosine fitting to C2, C3 and C4 as described above to the expression of oscillating genes in TC2, excluding genes with deviating mean expression in L4 as described above. We excluded C1, because amplitudes were sometimes difficult to call reliably. We used a fixed period of 7 h for C2-C3 and 8.5 h for C4 as determined by quantification of the oscillation period (Fig. 4C). We applied a linear regression using the function ‘lm’ of the package ‘stats’ in R to find the relationship between the amplitudes across different stages, i.e. the slope. The correlation coefficient, r, was determined using the ‘cor’ function (method=pearson) of the package ‘stats’ in R.

### Quantification of oscillation period

For a temporally resolved quantification of the oscillation period, we filtered the mean-normalized log_2_ transformed gene expression levels of oscillating genes, excluding L4 deviating genes (we selected TP5-TP39, because oscillations cease at ∼TP36 and the inclusion of 3 additional time points avoided edge effects) using a Butterworth filter (‘bwfilter’ function of the package ‘seewave’ (version 2.1.0) (Sueur et al., 2008) in R, to remove noise and trend-correct the data. The following command was used to perform the filtering: bwfilter(data, f = 1, n = 1, from = 0.1, to = 0.2, bandpass = TRUE, listen = FALSE, output = “matrix”). The bandpass frequency from 0.1 to 0.2 (corresponding to 10 hour and 5 hour period respectively) was selected based on the Fourier spectrum obtained after Fourier transform (‘fft’ function with standard parameters of the package ‘stats’). As an input for the Hilbert transform we used the butterworth-filtered gene expression. The ‘ifreq’ function (with standard parameters from the package ‘seewave’) was used to calculate the instantaneous phase and frequency based on the Hilbert transform. To determine the phase progression over time we unwrapped the instantaneous phase (ranging from 0 to 2π for each oscillation) using the ‘unwrap’ function of the package ‘EMD’ (version 1.5.7) (Kim and Oh, 2018) in R. To avoid edge effects, we removed the first 4 data points (TP5-TP8) and last 3 data points (TPTP37-TP39) of the unwrapped phase (retaining TP9-TP36). The angular velocity is defined as the rate of phase change, which we calculated by taking the derivative of the unwrapped phase. The instantaneous period was determined by 2π/angular velocity and was plotted for each gene individually and as mean in a density plot. The mean of the instantaneous period over all oscillating genes was used to reconstruct a ‘global’ oscillation by taking the following command: sin(cumsum(mean angular velocity)) and plotted together with a 7h-period oscillation and the mean normalized expression of a representative gene, *col-147*.

### Correlation analyses of RNAseq data

Log_2_-transformed data was filtered for oscillating genes and then plotted in a correlation matrix using the R command cor(data, method=”pearson”). The correlation line plots represent the correlations of selected time points to the fused full developmental time course (Fig. S5) and are specified in the line plot.

To reveal the highest correlations for a selected time point, we analyzed the correlation line of this time point between TP7 and TP36 (the time in which oscillations occur) using a spline analysis from Scipy (Jones et al., 2001) in python (“from scipy.interpolate import InterpolatedUnivariateSpline” with k=4) and stored the spline as variable “spline”. We identified peaks of the correlation line by finding the zeros of the derivative of the spline (cr_points = spline.derivative().roots()). The highest correlations of the respective correlation line were thus the value of the spline at the time point where the spline derivative was zero and the value was above the mean of the correlation line (cr_vals = spline(cr_pts) followed by pos_index = np.argwhere(cr_vals>np.mean(data.iloc[i])) and peak_val = cr_vals[pos_index]). Thus, we identified the correlation of particular time points (e.g. TP14–TP19) with their corresponding time points in the next oscillation cycle. Thereby, we were able to identify cycle time points as described in the results section. We defined the first cycle time point, e.g. TP14 of cycle 2, as 0°, and the last unique one, TP19, as 300°. TP14 (0° of cycle 2) is also 360° of cycle 1. Note that a sampling interval of 1 hour means that a TP in one cycle may correlate equally well to two adjacent TPs in another cycle, as seen for instance in the correlation of TP13 to TP26 and TP27. The spline interpolation places the peak of correlation in the middle of these time points at ∼TP26.5. The spline analysis thus annotates cycle points correctly even in C4 which has an extended period.

We performed correlation analyses without mean normalization of expression data, hence correlation values cannot be negative but remain between 0 and 1. We made this decision because a correlation analysis using mean-centered data, where correlations can vary between −1 and +1, requires specific assumptions on which time points to include or exclude for mean normalization, and because it is sensitive to gene expression trends. However, we confirmed, as a proof of principle, the expected negative correlation of time points that are in antiphase when using mean-centered data (Fig. S9) using all oscillating genes in TC3 (n = 3680).

### GO-term analysis

GO-term analysis was performed using the GO biological process complete option (GO ontology database, release 2019-02-02) from the online tool PANTHER (“PANTHER Classification System,” n.d.) (overrepresentation test, release 2019-03-08, standard settings).

### Tissue specific analysis

In order to reveal if particular tissues are enriched in oscillating genes, we used single cell sequencing data from (Cao et al., 2017). In particular, we used Supplementary “Table S6: Differential expression test results for the identification of tissue-enriched genes” where each gene’s highest and second highest tissue expression and the ratio of is reported. We selected tissue specific genes based on a ratio > 5 and a qvalue < 0.05 (these criteria reduced the number of genes to investigate). Using this list of genes we calculated the percentage of tissues present in both, all genes and oscillating genes using the function “Counter” from “collections” in python (*labels, values = zip(*Counter(tissue_info_thr[“max.tissue”]).items())*). In order to obtain the enrichment of tissues, we divided the percentage of tissue X among oscillating genes in the tissue enriched dataset by the percentage of tissue X among all genes in the tissue enriched data set and plotted the resulting values. The list of tissue specific oscillating genes was further used to investigate the peak phases within one tissue by plotting a density plot of the peak phase (from Fig. 1) for every tissue. As we lack data below 0 degree and above 360 degree, density values at these borders are distorted as the density is calculated over a moving window. Since we are confronted with cyclical data and thus 0 degree corresponds to 360 degree, we added and subtracted 360 degree to each phase value, thus creating data that ranged from −360 degree to 720 degree which allowed us to plot the correct density at the borders 0 and 360 degree. We used python (pandas) to plot this data using the following command:

data_tissue [“Phase”].plot(kind=“kde”, linewidth=5, alpha=0.5, bw=0.1)

### Identification of first gene expression peaks in L1 larvae

To identify the first peak of oscillating genes, we used a spline analysis in Python (“from scipy.interpolate import InterpolatedUnivariateSpline”) from TP3 – TP13. We chose these time points to remove false positives in the beginning due to slightly higher noise for the first 2 time points as well as not to identify the second peak which occurred at ≥TP14 for some very early genes. The function used was “InterpolatedUnivariateSpline” with k=4. After constructing the spline, we identified the zeros of the derivative and chose the time point value with the highest expression value and a zero derivative as the first peak time point.

### Embryonic gene expression time course

Embryonic gene expression data was obtained from (Hashimshony et al., 2015), and represented precisely staged single embryos at 10 min intervals from the 4-cell stage up to muscle movement and every 10-70 min thereafter until 830 minutes. We obtained the gene count data from the Gene Expression Omnibus data base under the accession number GSE50548, for which sequencing reads were mapped to WBCel215 genome and counted against WS230 annotation. We normalized the gene counts to the total mapped library size per sample, added a pseudocount of 8, and log_2_-transformed the data. We selected genes according to the larval oscillating gene annotation, with L4 deviating genes excluded, and plotted their embryonic expression patterns according to peak phase in larvae. The embryonic time course was correlated to the fused larval time course (TC3) using the ‘cor’ function (method=’pearson’) of the package ‘stats’ in R (Fig. S7A). Correlation line plots were generated by plotting the correlation coefficients for each embryonic time point over larval time. To identify the peaks of the correlation lines with a resolution higher than the sampling frequency, we interpolated the correlation lines using the ‘spline’ function (n=240, method=’fmm’) of the package ‘stats’ in R. To call the peaks of the interpolated correlation lines, we applied the ‘findpeaks’ function (with nups=5, ndowns=5) of the package ‘pracma’ on the time points on the interpolated time points 10-185, that cover the four cycles. To find the embryonic time point at which oscillations initiate, we plotted the larval TP in cycle 2 at which the correlation peak occurred over embryonic time (Fig. S7B) and determined the intersection of the two linear fits, using the ‘solve’ function of the package ‘Matrix’ (version 1.2-15) (Bates and Maechler, 2018) and the ‘lm’ function of the package ‘stats’ in R respectively. To determine the 95%-CI of the x-coordinate of the intersect, the standard error of the slope a and the intercept b of the two linear fits was propagated using Taylor expansion in the ‘propagate’ function (expr = expression((b1-b2)/(a2-a1)), ntype = “stat”,do.sim = FALSE, alpha=0.05) from the package ‘propagate’ in R. The pairwise correlation map was generated with the ‘aheatmap’ function of the package ‘NMF’ (version 0.21.0) (Gaujoux and Seoighe, 2010) and the 3D plot was generated with the ‘3Dscatter’ function of the package ‘plot3D’ (version 1.1.1) (Soetaert, 2017) in R.

### Time-lapse imaging of single animals

Single worm imaging was done by adapting a previous protocol (Turek et al., 2015), and is similar to the method reported in (Gritti et al., 2016). Specifically, we replaced the previous 3.5- cm dishes with a “sandwich-like” system: The bottom consisted of a glass cover slip onto which two silicone isolators (GRACE Bio-Labs, SKU: 666103) with a hole in the middle were placed on top of each other and glued onto the glass cover slip. We then placed single eggs inside the single OP50 containing chambers, which were made of 4.5% agarose in S-basal. The chambers including worms were then flipped 180 degree and placed onto the glass cover slip with the silicone isolators, so that worms faced the cover slip. Low melt agarose (3% in S-basal) was used to seal the agarose with the chambers to prevent drying out or drifts of the agarose chambers during imaging. The sandwich-like system was then covered with a glass slide on the top of the silicone isolators to close the system.

We used a 2x sCMOS camera model (T2) CSU_W1 Yokogawa microscope with 20x air objective, NA = 0.8 in combination with a 50µm disk unit to obtain images of single worms. For a high throughput, we motorized the stage positioning and the exchange between confocal and brightfield. We used a red LED light to combine brightfield with fluorescence without closing the shutter. Additionally, we used a motorized z-drive with 2 µm step size and 23 images per z-stack. The 488nm laser power for GFP imaging was set to 70% and a binning of 2 was used.

To facilitate detection of transgene expression and oscillation, we generated reporters using the promoters of genes that exhibited high transcript levels and amplitudes, and where GFP was concentrated in the nucleus and destabilized through fusion to PEST::H2B (see strain list above). We placed embryos into chambers containing food (concentrated bacteria HT115 with L4440 vector) and imaged every worm with a z-stack in time intervals of 10 min during larval development in a room kept at ∼21°C, using a double camera setting to acquire brightfield images in parallel with the fluorescent images. We exploited the availability of matching fluorescent and brightfield images to identify worms by machine learning. After identification, we flattened the worm at each time point to a single pixel line and stacked all time points from left to right, resulting in one kymograph image per worm. We then plotted background-subtracted GFP intensity values from the time of hatch (t = 0 h), which we identified by visual inspection of the brightfield images as the first time point when the worm exited the egg shell.

Time lapse images were analyzed using a customized KNIME workflow (Supplemental File 1). We analyzed every worm over time using the same algorithm. First, we identified the brightest focal planes per time point by calculating the mean intensity from all focal planes per time point and selecting the focal planes that had a higher intensity than the mean. Then we maximum-projected the GFP images over Z per time point and blurred the DIC image and also max projected over Z (blurring the DIC improved the machine learning process later on). All images per worm over time were analyzed by Ilastik machine learning in order to identify the worm in the image. The probability map from Ilastik was used to select a threshold that selected worms of a particular experiment best. (The threshold might change slightly as DIC images can look slightly different due to differences in the sample prep amongst experiments.) Using a customized ImageJ plugin, we straightened the worm. The straightened GFP worm image was then max projected over Y which resulted in a single pixel line representing the GFP intensities in a worm and after stacking up all the single pixel lines in Y direction, we obtained the kymographs. In order to remove noise coming from the head and tail regions of the worm due to inaccuracy of the machine learning, we measured mean GFP intensities per time point ranging from 20% until 80% of the worms anterior – posterior axis. For background subtraction we exploited the fact that only the nuclei were GFP positive and thus subtracted the minimum intensity value between GFP nuclei from their intensity values.

After the KNIME workflow, we imported the measured GFP intensities into Python and analyzed the traces using a butterworth filter and Hilbert transform analysis (both from Scipy (Jones et al., 2001)). We used the butterworth bandpass filter using b, a = butter(order =1, [low,high], btype=”band”) with low=1/14 and high=1/5, corresponding to 14 hour and 5 hour periods respectively. We then filtered using filtfilt(b, a, data, padtype=’constant’) to linearly filter backwards and forwards.

For individual time points where the worm could not be identified by the Ilastik machine learning algorithm, we linearly interpolated (using interpolation from pandas (McKinney, 2010)) using “pandas.series.interpolate(method = ‘linear’, axis =0, limit = 60, limit_direction = ‘backward’”, between the neighboring time points to obtain a continuous time series needed for the Hilbert transform analysis. Using Hilbert transform, we extracted the phase of the oscillating traces for each time point and specifically investigated the phase at molt entry and molt exit for our different reporter strains.

In order to determine time points in which worms are in lethargus, we investigated pumping behavior. As the z-stack of an individual time point gives a short representation of a moving worm, it is possible to determine whether animals pump (feeding, corresponds to intermolt) or not (lethargus / molt). Additionally to the pumping behavior, we used two further requirements that needed to be true in order to assign the lethargus time span: First, worms needed to be quiescent (not moving, and straight line) and second, a cuticle needed to be shed at the end of lethargus. Usually worms start pumping one to two time points before they shed the cuticle. This analysis was done manually with the software ImageJ, and results were recorded in an excel file, where for every time point, the worms’ behavior was denoted as 1 for pumping and as 0 for non-pumping.

To determine a possible connection between oscillations and development, we applied error propagation, assuming normal distribution of the measured phases and larval stage durations. Thereby, we exploited the inherent variation of the oscillation periods and developmental rates among worms, rather than experimental perturbation, to probe for such a connection. We define the phase *θ* at either molt exit or entry as 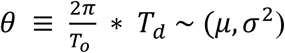 with 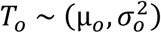 being the period of oscillation and 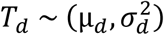 the intermolt duration (for phase at molt entry) or larval stage duration (for phase at molt exit), resulting in a phase with mean *μ* and a standard deviation *σ*. Should the two processes be coupled as in scenario 2, we would expect

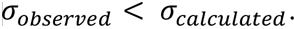

To calculate the phase at molt entry and molt exit with error propagation we used the “uncertainties” package (Lebigot, n.d.) in python. The larval stage duration as well as intermolt duration and period were treated as ufloat numbers, representing the distributions coming from our measurement (e.g. 7.5 +/- 0.2). These distributions were then used to calculate the expected phase at molt entry (using the intermolt duration) and molt exit (using the larval stage duration) using: 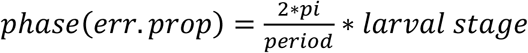. This resulted in the phase being represented by an ufloat number and thus a distribution which we used for plotting after normalizing for the mean to compare the variation of the data.

## References

1. Abraham, U., Granada, A.E., Westermark, P.O., Heine, M., Kramer, A., Herzel, H., 2010. Coupling governs entrainment range of circadian clocks. Mol Syst Biol 6, 438. https://doi.org/10.1038/msb.2010.92

2. Au, K.F., Jiang, H., Lin, L., Xing, Y., Wong, W.H., 2010. Detection of splice junctions from paired-end RNA-seq data by SpliceMap. Nucleic Acids Res 38, 4570–4578. https://doi.org/10.1093/nar/gkq211

3. Bates, D., Maechler, M., 2018. Matrix: Sparse and Dense Matrix Classes and Methods.

4. Baugh, L.R., 2013. To grow or not to grow: nutritional control of development during Caenorhabditis elegans L1 arrest. Genetics 194, 539–555. https://doi.org/10.1534/genetics.113.150847

5. Cao, J., Packer, J.S., Ramani, V., Cusanovich, D.A., Huynh, C., Daza, R., Qiu, X., Lee, C., Furlan, S.N., Steemers, F.J., Adey, A., Waterston, R.H., Trapnell, C., Shendure, J., 2017. Comprehensive single-cell transcriptional profiling of a multicellular organism. Science 357, 661–667. https://doi.org/10.1126/science.aam8940

6. Conrad, E., Mayo, A.E., Ninfa, A.J., Forger, D.B., 2008. Rate constants rather than biochemical mechanism determine behaviour of genetic clocks. J R Soc Interface 5, S9–S15. https://doi.org/10.1098/rsif.2008.0046.focus

7. Edgar, R., Domrachev, M., Lash, A.E., 2002. Gene Expression Omnibus: NCBI gene expression and hybridization array data repository. Nucleic Acids Res 30, 207–210. https://doi.org/10.1093/nar/30.1.207

8. Forger, D.B., 2017. Biological Clocks, Rhythms, and Oscillations: The Theory of Biological Timekeeping. The MIT Press, Cambridge, Massachusetts.

9. Frand, A.R., Russel, S., Ruvkun, G., 2005. Functional Genomic Analysis of C. elegans Molting. PLOS Biology 3, e312. https://doi.org/10.1371/journal.pbio.0030312

10. Frøkjær-Jensen, C., Davis, M.W., Ailion, M., Jorgensen, E.M., 2012. Improved Mos1-mediated transgenesis in C. elegans. Nat. Methods 9, 117–118. https://doi.org/10.1038/nmeth.1865

11. Frøkjaer-Jensen, C., Davis, M.W., Hopkins, C.E., Newman, B.J., Thummel, J.M., Olesen, S.-P., Grunnet, M., Jorgensen, E.M., 2008. Single-copy insertion of transgenes in Caenorhabditis elegans. Nat. Genet. 40, 1375–1383. https://doi.org/10.1038/ng.248

12. Gaidatzis, D., Lerch, A., Hahne, F., Stadler, M.B., 2015. QuasR: quantification and annotation of short reads in R. Bioinformatics 31, 1130–1132. https://doi.org/10.1093/bioinformatics/btu781

13. Gaujoux, R., Seoighe, C., 2010. A flexible R package for nonnegative matrix factorization. BMC Bioinformatics 11, 367. https://doi.org/10.1186/1471-2105-11-367

14. Gibson, D.G., Young, L., Chuang, R.-Y., Venter, J.C., Hutchison, C.A., Smith, H.O., 2009. Enzymatic assembly of DNA molecules up to several hundred kilobases. Nat. Methods 6, 343–345. https://doi.org/10.1038/nmeth.1318

15. Gritti, N., Kienle, S., Filina, O., van Zon, J.S., 2016. Long-term time-lapse microscopy of *C. elegans* post-embryonic development. Nature Communications 7, 12500. https://doi.org/10.1038/ncomms12500

16. Grün, D., Kirchner, M., Thierfelder, N., Stoeckius, M., Selbach, M., Rajewsky, N., 2014. Conservation of mRNA and protein expression during development of C. elegans. Cell Rep 6, 565–577. https://doi.org/10.1016/j.celrep.2014.01.001

17. Guantes, R., Poyatos, J.F., 2006. Dynamical principles of two-component genetic oscillators. PLoS Comput. Biol. 2, e30. https://doi.org/10.1371/journal.pcbi.0020030

18. Hall, D.H., Herndon, L.A., Altun, Z., 2017. Introduction to C. elegans Embryo Anatomy. In WormAtlas.

19. Hashimshony, T., Feder, M., Levin, M., Hall, B.K., Yanai, I., 2015. Spatiotemporal transcriptomics reveals the evolutionary history of the endoderm germ layer. Nature 519, 219–222. https://doi.org/10.1038/nature13996

20. Hendriks, G.-J., Gaidatzis, D., Aeschimann, F., Großhans, H., 2014. Extensive oscillatory gene expression during C. elegans larval development. Mol. Cell 53, 380–392. https://doi.org/10.1016/j.molcel.2013.12.013

21. Izhikevich, E.M., 2000. Neural excitability, spiking and bursting. Int. J. Bifurcation Chaos 10, 1171–1266. https://doi.org/10.1142/S0218127400000840

22. Jensen, P.B., Pedersen, L., Krishna, S., Jensen, M.H., 2010. A Wnt Oscillator Model for Somitogenesis. Biophys J 98, 943–950. https://doi.org/10.1016/j.bpj.2009.11.039

23. Jones, E., Oliphant, T., Peterson, P., 2001. SciPy: Open Source Scientific Tools for Python.

24. Kim, D., Oh, H.-S., 2018. EMD: Empirical Mode Decomposition and Hilbert Spectral Analysis.

25. Kim, D. hyun, Grün, D., van Oudenaarden, A., 2013. Dampening of expression oscillations by synchronous regulation of a microRNA and its target. Nat. Genet. 45, 1337–1344. https://doi.org/10.1038/ng.2763

26. Koike, N., Yoo, S.-H., Huang, H.-C., Kumar, V., Lee, C., Kim, T.-K., Takahashi, J.S., 2012. Transcriptional Architecture and Chromatin Landscape of the Core Circadian Clock in Mammals. Science 338, 349–354. https://doi.org/10.1126/science.1226339

27. Korenčič, A., Košir, R., Bordyugov, G., Lehmann, R., Rozman, D., Herzel, H., 2014. Timing of circadian genes in mammalian tissues. Sci Rep 4, 1–9. https://doi.org/10.1038/srep05782

28. Lagido, C., Pettitt, J., Flett, A., Glover, L.A., 2008. Bridging the phenotypic gap: Real-time assessment of mitochondrial function and metabolism of the nematode Caenorhabditis elegans. BMC Physiology 8, 7. https://doi.org/10.1186/1472-6793-8-7

29. Lažetić, V., Fay, D.S., 2017. Molting in C. elegans. Worm 6, e1330246. https://doi.org/10.1080/21624054.2017.1330246

30. Lebigot, E.O., n.d. Uncertainties: a Python package for calculations with uncertainties.

31. McKinney, W., 2010. Data Structures for Statistical Computing in Python, in: Proceedings of the 9th Python in Science Conference. pp. 51–56.

32. Mönke, G., Cristiano, E., Finzel, A., Friedrich, D., Herzel, H., Falcke, M., Loewer, A., 2017. Excitability in the p53 network mediates robust signaling with tunable activation thresholds in single cells. Scientific Reports 7, 46571. https://doi.org/10.1038/srep46571

33. Moreno-Risueno, M.A., Van Norman, J.M., Moreno, A., Zhang, J., Ahnert, S.E., Benfey, P.N., 2010. Oscillating gene expression determines competence for periodic Arabidopsis root branching. Science 329, 1306–1311. https://doi.org/10.1126/science.1191937

34. Niwa, R., Hada, K., Moliyama, K., Ohniwa, R.L., Tan, Y.-M., Olsson-Carter, K., Chi, W., Reinke, V., Slack, F.J., 2009. C. elegans sym-1 is a downstream target of the hunchback-like-1 developmental timing transcription factor. Cell Cycle 8, 4147–4154. https://doi.org/10.4161/cc.8.24.10292

35. Oates, A.C., Morelli, L.G., Ares, S., 2012. Patterning embryos with oscillations: structure, function and dynamics of the vertebrate segmentation clock. Development 139, 625–639. https://doi.org/10.1242/dev.063735

36. Olmedo, M., Geibel, M., Artal-Sanz, M., Merrow, M., 2015. A High-Throughput Method for the Analysis of Larval Developmental Phenotypes in Caenorhabditis elegans. Genetics 201, 443–448. https://doi.org/10.1534/genetics.115.179242

37. Panda, S., Hogenesch, J.B., Kay, S.A., 2002. Circadian rhythms from flies to human. Nature 417, 329–335. https://doi.org/10.1038/417329a

38. PANTHER Classification System, n.d.

39. Pikovsky, A., Rosenblum, M., Kurths, J., 2001. Synchronization: A universal concept in nonlinear sciences. Cambridge University Press, Cambridge. https://doi.org/10.1017/CBO9780511755743

40. Purcell, O., Savery, N.J., Grierson, C.S., di Bernardo, M., 2010. A comparative analysis of synthetic genetic oscillators. J R Soc Interface 7, 1503–1524. https://doi.org/10.1098/rsif.2010.0183

41. R Core Team, n.d. R: A language and environment for statistical computing.

42. Rensing, L., Meyer-Grahle, U., Ruoff, P., 2001. Biological timing and the clock metaphor: oscillatory and hourglass mechanisms. Chronobiol. Int. 18, 329–369.

43. Riedel-Kruse, I.H., Müller, C., Oates, A.C., 2007. Synchrony dynamics during initiation, failure, and rescue of the segmentation clock. Science 317, 1911–1915. https://doi.org/10.1126/science.1142538

44. Ruaud, A.-F., Bessereau, J.-L., 2006. Activation of nicotinic receptors uncouples a developmental timer from the molting timer in C. elegans. Development 133, 2211–2222. https://doi.org/10.1242/dev.02392

45. Saggio, M.L., Spiegler, A., Bernard, C., Jirsa, V.K., 2017. Fast–Slow Bursters in the Unfolding of a High Codimension Singularity and the Ultra-slow Transitions of Classes. J Math Neurosci 7. https://doi.org/10.1186/s13408-017-0050-8

46. Salvi, J.D., Ó Maoiléidigh, D., Hudspeth, A.J., 2016. Identification of Bifurcations from Observations of Noisy Biological Oscillators. Biophys. J. 111, 798–812. https://doi.org/10.1016/j.bpj.2016.07.027

47. Schindler, A.J., Baugh, L.R., Sherwood, D.R., 2014. Identification of late larval stage developmental checkpoints in Caenorhabditis elegans regulated by insulin/IGF and steroid hormone signaling pathways. PLoS Genet. 10, e1004426. https://doi.org/10.1371/journal.pgen.1004426

48. Shih, N.P., François, P., Delaune, E.A., Amacher, S.L., 2015. Dynamics of the slowing segmentation clock reveal alternating two-segment periodicity. Development 142, 1785–1793. https://doi.org/10.1242/dev.119057

49. Soetaert, K., 2017. plot3D: Plotting Multi-Dimensional Data.

50. Soroldoni, D., Jörg, D.J., Morelli, L.G., Richmond, D.L., Schindelin, J., Jülicher, F., Oates, A.C., 2014. Genetic oscillations. A Doppler effect in embryonic pattern formation. Science 345, 222–225. https://doi.org/10.1126/science.1253089

51. Spiess, A.-N., 2018. propagate: Propagation of Uncertainty.

52. Stiernagle, Theresa, n.d. Maintenance of C. elegans (February 11, 2006), WormBook, ed. The C. elegans Research Community, WormBook, doi/10.1895/wormbook.1.101.1, http://www.wormbook.org.

53. Strogatz, S.H., 2015. Nonlinear Dynamics and Chaos with Student Solutions Manual: Nonlinear Dynamics and Chaos: With applications To Physics, Biology, Chemistry, And Engineering (Studies in Nonlinearity), 2nd ed. Westview Press, Boulder, CO.

54. Sueur, J., Aubin, T., Simonis, C., 2008. Seewave, a Free Modular Tool for Sound Analysis and Synthesis. Bioacoustics 18, 213–226. https://doi.org/10.1080/09524622.2008.9753600

55. Sulston, J.E., Schierenberg, E., White, J.G., Thomson, J.N., 1983. The embryonic cell lineage of the nematode Caenorhabditis elegans. Dev. Biol. 100, 64–119.

56. Tsai, T.Y.-C., Choi, Y.S., Ma, W., Pomerening, J.R., Tang, C., Ferrell, J.E., 2008. Robust, Tunable Biological Oscillations from Interlinked Positive and Negative Feedback Loops. Science 321, 126–129. https://doi.org/10.1126/science.1156951

57. Turek, M., Besseling, J., Bringmann, H., 2015. Agarose Microchambers for Long-term Calcium Imaging of Caenorhabditis elegans. J Vis Exp e52742. https://doi.org/10.3791/52742

58. Uriu, K., 2016. Genetic oscillators in development. Development, Growth & Differentiation 58, 16–30. https://doi.org/10.1111/dgd.12262

59. Vuong-Brender, T.T.K., Ben Amar, M., Pontabry, J., Labouesse, M., 2017. The interplay of stiffness and force anisotropies drives embryo elongation. eLife 6, e23866. https://doi.org/10.7554/eLife.23866

60. Webb, A.B., Lengyel, I.M., Jörg, D.J., Valentin, G., Jülicher, F., Morelli, L.G., Oates, A.C., 2016. Persistence, period and precision of autonomous cellular oscillators from the zebrafish segmentation clock. eLife 5, e08438. https://doi.org/10.7554/eLife.08438

61. Zhang, R., Lahens, N.F., Ballance, H.I., Hughes, M.E., Hogenesch, J.B., 2014. A circadian gene expression atlas in mammals: implications for biology and medicine. Proc. Natl. Acad. Sci. U.S.A. 111, 16219–16224. https://doi.org/10.1073/pnas.1408886111

